# Induction of cervical dysfunction associated with preterm birth by IL-1 and dysbiotic microbiome revealed in human endocervix chips

**DOI:** 10.1101/2025.05.01.651107

**Authors:** Anna Stejskalova, Karina Calderon, David B. Chou, Aakanksha Gulati, Jessica F. Feitor, Zohreh Izadifar, Ola Gutzeit, Yasmine Bouchibti, Siyu Chen, Roberto Plebani, Justin Cotton, Jasmine Hughes, Thomas Ferrante, Bogdan Budnik, Girija Goyal, Abidemi Junaid, Carlito B. Lebrilla, Donald E. Ingber

## Abstract

Cervical dysfunction, a major contributor to preterm labor and neonatal mortality, remains poorly understood due to the absence of physiologically relevant human models. Here, we show that a microfluidic human Endocervix Chip lined by primary endocervical epithelium interfaced with stromal cells and cultured under pregnancy-like hormonal conditions recapitulates key aspects of the biology of the endocervix, including formation of a mucus plug-like structure with antimicrobial properties. Culturing a dysbiotic cervico-vaginal microbiome on-chip increased secretion of pro-inflammatory cytokines observed in patients with preterm labor and enhanced production of matrix metalloproteinases (MMPs) that degrade stromal extracellular matrix (ECM). Perfusion with inflammatory cytokines at clinically relevant concentrations altered cervical mucus composition, upregulated prostaglandin-endoperoxide synthase 2 expression, increased MMP secretion, and reduced collagen production, which together drive dissolution of the stromal ECM and promote cervical ripening. Addition of circulating peripheral blood mononuclear cells (PBMCs) amplified these effects. Administration of a clinically approved drug for prevention of preterm labor that the Food and Drug Administration (FDA) recently deemed ineffective was also found to be inactive in the chip, while an approved therapeutic antagonist of the IL-1 receptor successfully protected against cervical dysfunction in this model. These findings demonstrate that IL-1 acts directly on human cervical tissues to promote changes associated with initiation of labor and that primary human endocervix chips may represent a useful preclinical model for studies on cervical dysfunction associated with preterm birth.

**One Sentence Summary:** Human endocervix chip modeling of cervical dysfunction suggests that IL-1 receptor antagonists may prevent preterm labor.

## INTRODUCTION

Preterm birth (birth before the 37^th^ week of gestation) affects more than 13 million infants a year (*1*), and it is the second leading cause of early childhood mortality and disability (*2*) affecting the lifelong well-being of individuals, families, and communities. Despite the prevalence and life-long consequences of prematurity, there are currently no effective therapeutic options to prevent preterm birth. The risk of preterm labor is also influenced by the commensal cervico-vaginal microbiome with a *Lactobacillus crispatus (L. crispatus)* dominated microbiome being associated with health and full term labor (*3*) while the presence of *L. crispatus*-depleted non-optimal (dysbiotic) communities correlates with an increased risk of cervical shortening and preterm labor (*3*, *4*). It is, however, unclear how a dysbiotic cervico-vaginal microbiome increases this risk and why only some women with this condition deliver prematurely (*5*, *6*).

Protecting a developing fetus from pathogenic microbes and associated inflammation throughout gestation is essential, but also challenging clinically because it requires a balancing act, given that labor itself is considered to be an inflammatory process (*5*). However, nature has evolved a way to protect the fetus from potential pathogenic microbes in the form of a specialized mucus plug rich in antimicrobial peptides and mucins that accumulates in the cervical lumen, which enhances epithelial mucosal defenses(*6*). The cervix also contracts and mechanically holds the cervical lumen tightly closed throughout pregnancy, providing a physical barrier (*7*); then just prior to delivery, the cervix undergoes a ripening process in which it softens, shortens, and subsequently dilates enabling passage of the fetus through the birth canal (*8*). The cervix can also dysfunction and premature cervical shortening (cervical length < 25 mm before the 24^th^ week of pregnancy) is currently the only clinically recognized early risk factor for preterm labor that is screened for routinely.

Unfortunately, studying both physiological and premature human labor, including premature loosening of the mucus plug and abnormal cervical ripening, has been technically and ethically challenging because virtually all animal models except for non-human primates have a different mechanism of labor (*9*). In addition, host-microbiome interactions are highly species-specific (*10*). Consequently, there has been little progress in the development of new therapies for preterm labor since the 1960s (*11*). Recent advances in fetal-maternal cell culture protocols (*12–14*) and advances in microphysiological systems (*15*) have offered new hope to bridge this research gap, however, there is still a paucity of human relevant models.

Here, we leveraged organ-on-a-chip (Organ Chip) microfluidic culture technology to develop primary human Endocervix Chips lined by hysterectomy-derived human endocervical epithelial cells interfaced with donor tissue-matched stromal fibroblasts, which were used with or without peripheral blood mononuclear cells (PBMCs) to model physiological and pathophysiological states of the human cervix *in vitro*. The Endocervix Chips were employed to explore how human-relevant cervicovaginal microbiome and related inflammation contribute to cervical dysfunction under pregnancy-like hormonal conditions, as well as how progestin, non-steroidal anti-inflammatory drugs (NSAIDs), and cytokine-suppressive anti-inflammatory drugs (CSAIDs) including an IL-1 antagonist modulate cervix structure and function. Past work revealed that IL-1 plays an important role in control of preterm labor by stimulating uterine contractions (*16–18*), however, its direct effects on cervical tissues have never been explored. Our results show that this Organ Chip model of the human endocervix represents a new preclinical model that can be used to study cervical dysfunction caused by dysbiotic microbiome and inflammation. The work also reveals that IL-1 acts directly on cervical tissue to induce changes that contribute to preterm labor and that IL-1 antagonists may offer a new means of therapeutic intervention.

## RESULTS

### Fabrication of Primary Human Cervix Chips

The cervix serves as a physical and biochemical barrier between the microbiome-rich vagina and uterus (*7*). It is composed of a lining epithelium and an underlying stroma that is divided into the ectocervix, which contains an outward facing epithelium on its lower end that directly interfaces with the vagina, and the endocervix that consists of a simple columnar epithelium that lines the internal surface of the birth canal at the lower end of the uterus. Given its location, the endocervix plays a more central role in maintaining a successful pregnancy. We recently described a protocol to create a 2-channel, microfluidic, human Cervix Chip lined with a mixed population of commercially available ecto– and endo-cervical epithelial cells interfaced with stromal cells (*19*). Here, we modified this protocol to fabricate a primary human Endocervix Chip lined with hysterectomy-derived endocervical epithelial cells cultured in one channel that are interfaced across a porous ECM-coated membrane with matched cervical stromal cells isolated from the same donor tissue (**Fig. 1A, fig. S1A-E**). Ectocervix Chips were also fabricated using a similar approach (**fig. S1A-E**), which we used as controls.

**Figure 1.**
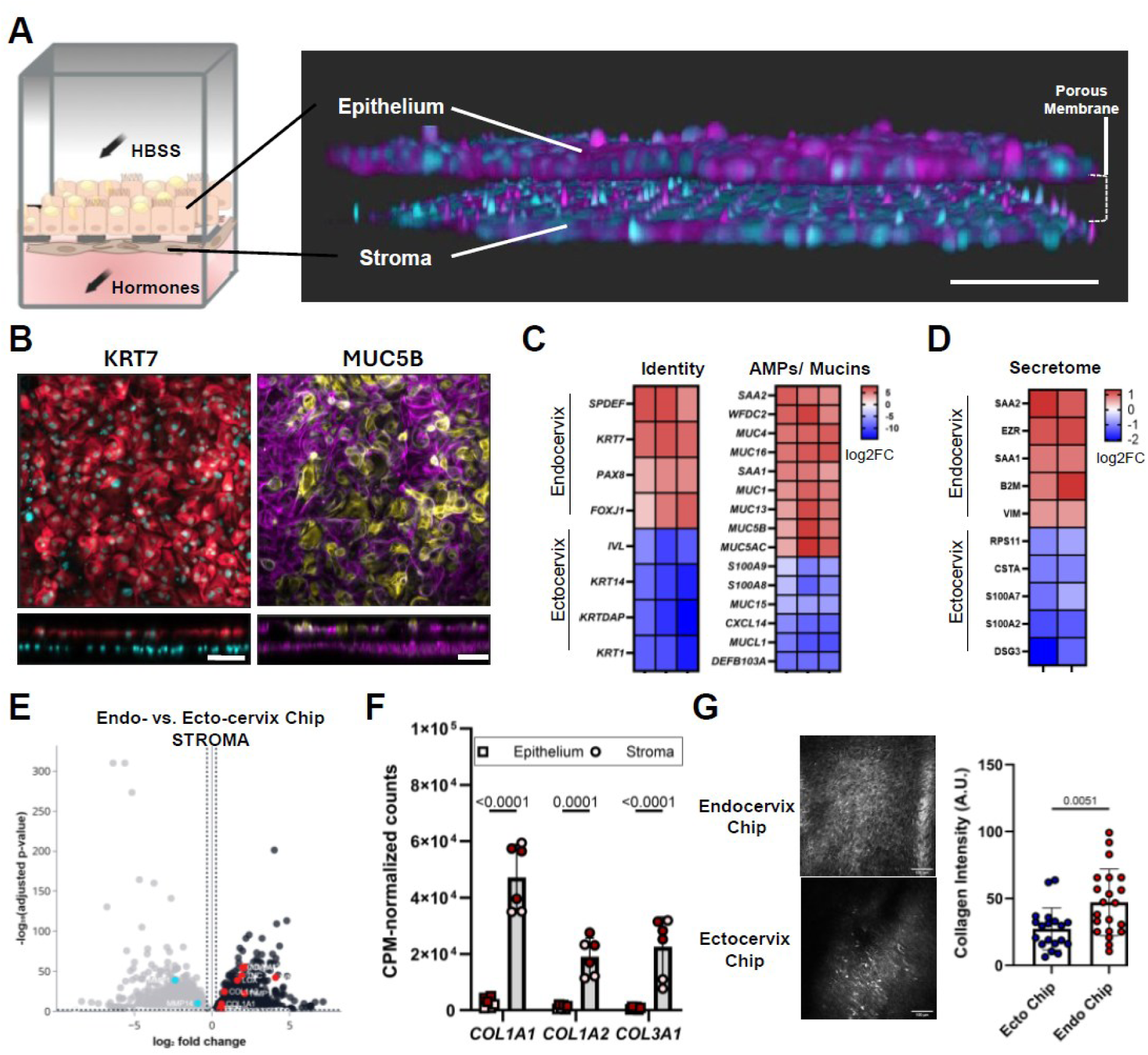
Human Endocervix Chip Development and Characterization. **A**) Schematic of the Endocervix Chip (left) and confocal reconstruction of a vertical cross section through an Endocervix Chip (right) showing the epithelium in the apical channel and stromal fibroblasts in the basal channel stained for F-actin (purple) and nuclei (blue) separated by a porous membrane that is not visible in this view (bar, 150 μm). **B**) Immunofluorescence microscopic images of the endocervical epithelium on-chip viewed from above stained for endocervical markers KRT7 (Left, red) and MUC5B (right, yellow) as well as F-actin (purple) and nuclei (blue). Bottom images show vertical cross sectional views of the same images (bar, 100 μm, left and 80 μm, right). **C**) Heatmap showing significantly upregulated and downregulated genes in the epithelium of Endocervix Chip versus Ectocervix Chip relating to endocervical versus ectocervical epithelial cells, antimicrobial peptides (AMPs) and mucins (log2FC = log2 fold change; N=3 endocervix donors, N=3-4 chips per donor). **D**) Heatmap showing significantly upregulated and downregulated secreted proteins (<30 kDa) in the apical effluents of Endocervix Chips versus Ectocervix Chips (log2FC = log2 fold change; N=2 donors, N=4 chips per group). **E**) Volcano plot comparing gene expression by stromal cells within donor matched Endocervix (black) and Ectocervix (grey) Chips showing log2 fold change of each gene on the x-axis and the –log10(adjusted p-value) on the y-axis, which demonstrate unique gene expression profiles for endocervical versus ectocervical stroma. Points represent genes with adjusted p-value ≤ 0.01 and a fold change that is either < 0.8 or >1.2. (N=4 chips per group). **F**) Bar graph showing gene expression levels expressed as counts per million (CPM)-normalized counts for the collagens *COL1A1*, *COL1A2,*and *COL3A1* (□ epithelium, apical lysate ○ stroma, basal lysate; N=2 human donors, 3 chips per donor, Mean ± SD, two-way ANOVA, multiple comparisons). **G**) Second harmonic microscopic images (left) showing fibrillar collagen in the basal channel of an Endocervix Chip versus Ectocervix Chip cultured in E2 containing medium (bar, 100μm) with results of quantification of signal intensity shown in a graph at the right (a.u., arbitrary units, N=3-4 chips, n=19-21 unique fields of view, Mean ± SD, *t*-test).

Freshly isolated epithelial cells and stromal cells were first expanded in conventional culture dishes in medium rich in Wnt3a, R-spondin-3 and Noggin (*13*, *14*). Immunofluorescence microscopic analysis confirmed that the primary human ectocervical epithelial cells expressed keratin 7 (KRT7), as endocervical epithelial cells do *in vivo* (*13*) (**fig. S2A).** In contrast, most of the primary stromal cells isolated from the same endocervical tissue expressed CD34 and vimentin (**fig. S2B-D**), which are markers of endocervical stromal fibroblasts, although a small fraction were likely smooth muscle cells as they were desmin-positive (**fig. S2B-D**). To create the Endocervix Chips, these cells were then enzymatically detached from the culture dish and cultured in parallel channels on opposite sides of a common ECM-coated porous membrane within a commercially available 2-channel Organ Chip microfluidic culture device (Emulate Inc.). After cell expansion for 5 days on-chip, the apical nutrient-rich expansion medium was replaced with Hank’s Balanced Saline Solution with calcium (HBSS), while differentiation medium containing estradiol (E2) was flowed continuously (40μl/h) through the basal channel to promote tissue differentiation. When cultured under these dynamic flow conditions, the endocervical epithelium formed a monolayer interfaced with a fibroblast-rich stroma on-chip, hence recapitulating the epithelial-stromal interface observed within the endocervix *in vivo* (**Fig. 1A).** Primary human Ectocervix Chips were created by replacing the endocervical cells with hysterectomy-derived ectocervical epithelial and stromal cells from the same donor.

When we compared epithelial production of KRT7 and the gel forming mucin MUC5B, we confirmed that both were only expressed in the differentiated Endocervix Chips (**Fig. 1B)** and not in the Ectocervix Chips (**fig. S3A),** which is consistent with expression patterns observed *in vivo* (*13*). Differential gene expression analysis of epithelia in the donor-matched Endocervix Chips versus Ectocervix Chips showed that each epithelium displayed a distinct pattern of tissue-specific gene expression (**fig. S3B**), which aligned with their known distinct biological roles in human pregnancy and reproduction. For example, epithelium in the Endocervix Chips expressed higher levels of the endocervical identity and lineage markers (e.g., *SPDEF*, *KRT7*, *PAX8*, *FOXJ1*) while genes commonly expressed by the ectocervical epithelium (e.g., *IVL*, *KRT14*, *KRTDAP*, *KRT1*) were more highly expressed in the Ectocervix Chips (**Fig. 1C**). Similarly, multiple endocervical mucin genes (e.g., *MUC1*, *MUC4*, *MUC5AC*, *MUC5B*, *MUC13*, *MUC16*) were more prominent in the Endocervix Chips while the Ectocervix Chips expressed significantly higher levels of the transmembrane mucins *MUCL1* and *MUC15.* Epithelial cells in both chips also expressed genes encoding antimicrobial peptides (AMPs) that are critical modulators of microbial niche composition across epithelia, with *SAA2* and *WFDC2* being more highly expressed by the endocervical epithelium while the ectocervical cells expressed *S100A8*, *S100A9*, *CXCL14* and *DEFB103A* (**Fig. 1C**).

Importantly, while vaginal secretions collected from patients cannot be used to distinguish between secretions from different parts of the female reproductive tract, we were able to address this challenge directly using engineered human Organ Chips. Unbiased analysis of proteins contained within apical chip outflows revealed several differentially expressed AMPs, including higher levels of serum amyloid A1 and A2, SAA1, and SAA2 in the Endocervix Chips, which are all proteins with antimicrobial properties (*20*), while the Ectocervix Chip secretome was enriched for S100A2 and psoriasin (S100A7) (**Fig. 1D, fig. S3C**).

These results are consistent with our transcriptomics data confirming that *SAA1* and *SAA2* are expressed at higher levels in endocervical epithelium (**fig. S3B).** Gene set enrichment analysis (GSEA) of this dataset also showed that the gene profiles of the Endocervix Chip epithelium were enriched for terms related to cellular extravasation, defense response to virus, inflammatory response and innate immune response, while processes related to cornification and epithelial cell differentiation dominated in the Ectocervix Chips (**fig. S3D**), again as expected from their physiological roles *in vivo*

Interestingly, the stromal compartment of the Ectocervix Chips also displayed a distinct pattern of gene expression (**Fig. 1E**) including higher expression of genes encoding proteins related to ECM breakdown, such as the matrix metalloproteinase 1 (*MMP1),* than stroma cells cultured in the Endocervix Chip that expressed higher levels of genes encoding multiple fibrillar collagens (*COL1A1*, *COL1A2*, *COL3A1),* decorin (*DCN*), elastin (*ELN*) and hyaluronic acid synthase 2 (*HAS2*), tissue inhibitor of metalloproteinase 1 (*TIMP1*), and the ECM crosslinking enzyme lysyl oxidase (*LOX*) (**Fig. 1E,F)**. Importantly, these changes at the level of gene expression correlated with stromal production of a thicker fibrillar collagenous matrix in the Endocervix Chip compared to the Ectocervix Chip, as visualized and quantified using second harmonic imaging (**Fig. 1G**). This is an important finding because accumulation and maintenance of an intact fibrillar collagenous ECM is a key feature of endocervix function during normal pregnancy. Moreover, dissolution of the ECM within the cervix is associated with onset of premature labor (*8*), and thus, it must be reliably reproduced in any *in vitro* study that focuses on cervical dysfunction.

### Mucus plug formation within Endocervix Chips

Animal models have provided insights into the structural and biochemical changes that occur in the cervix during pregnancy(*11*) but there is little information currently available about the mucus plug formed by the human endocervix, which protects against ascending infection (*21*). It is known that female hormones play a key role in modulating mucus plug formation as well as ECM turnover and immune responses in the cervix (*22*), and thus, that relevant hormones must be incorporated in any model used in reproductive biology research to study effects of pregnancy(*12*). We therefore mimicked pregnancy on-chip by administering a previously described mixture (*12*) of pregnancy-associated hormones (PAHs), which include E2, medroxyprogesterone acetate, human chorionic gonadotropin, human placental lactogen and prolactin, with concentrations adjusted to more closely reflect the early 3^rd^ trimester of pregnancy that is most relevant to preterm labor (*23*, *24*) (**table S1**).

The structural properties of cervical mucus are important during pregnancy because in addition to forming a thickened mucus plug that separates the epithelium from overlying bacteria, mucus flow enhances bacterial clearance. Importantly, we found that the Endocervix Chips deposit a thick mucus plug-like structure that fills the apical channel, as visualized using the lectins Jacalin and Wheat Germ Agglutinin (WGA) that stain for MUC5AC and MUC5B, respectively (*25*) (**Fig. 2A**). Interestingly, the Jacalin staining localized more closely to the epithelium while WGA staining appeared in an expanded fibrillar pattern throughout mucus-filled lumen. A similar pattern of mucin organization with MUC5AC creating sheets and MUC5B forming strands was previously seen in the lung airways (*25*). Thus, the spatial arrangement of these two co-expressed gel-forming mucins appears to be shared between these different types of epithelia that also share a common function: maintaining optimal microbial clearance.

**Figure 2.**
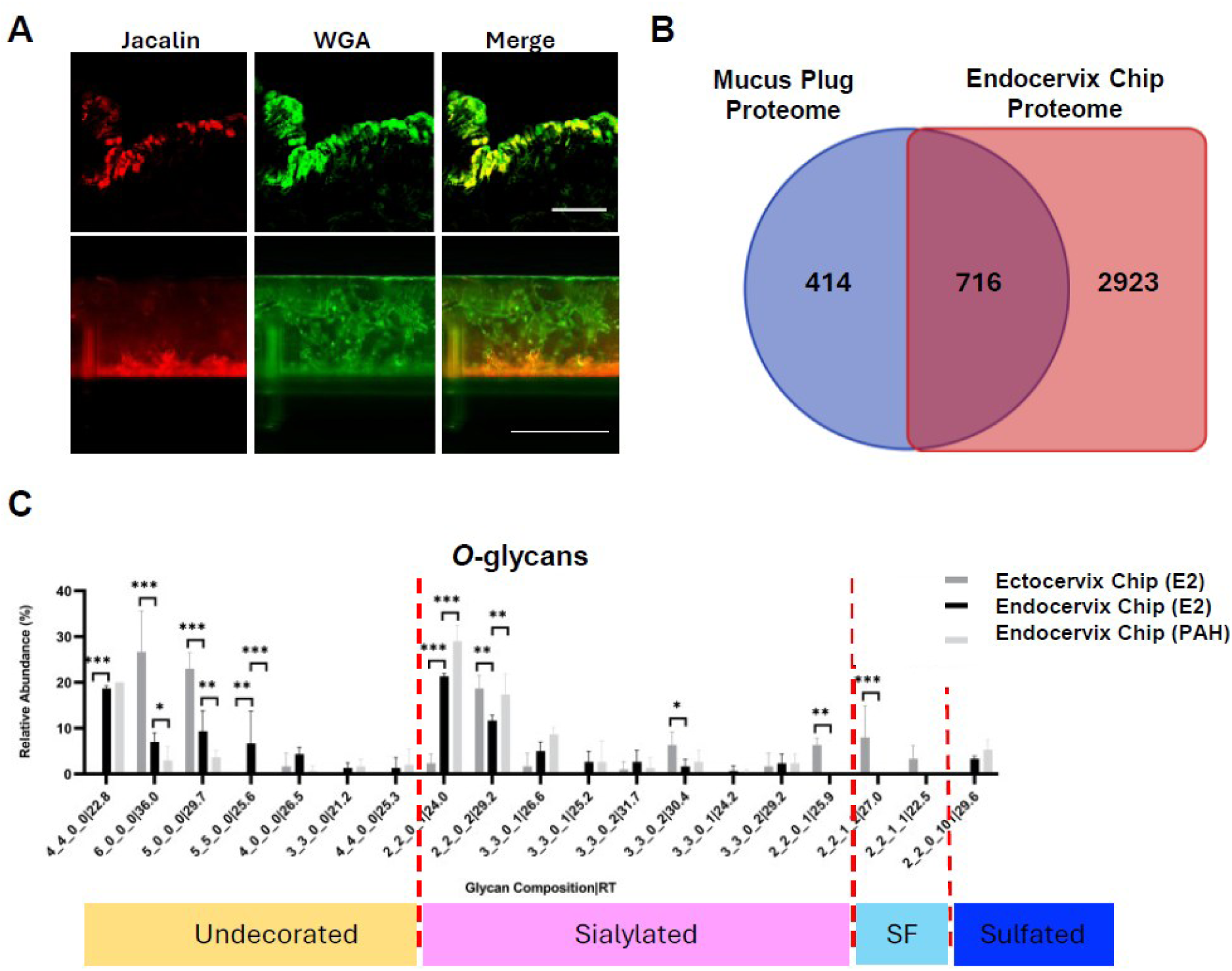
Mucus Characterization in Endocervix Chips. **A**) Immunofluorescence microscopic images of histological sections through snap-frozen human endocervix tissue (top) and side view images of a human Endocervix Chip (bottom) showing the mucus layer visualized with Jacalin (red) and Wheat Germ Agglutinin (WGA, green) (bar, 25 μm, tissue; 1 mm, chip). **B**) Venn diagram showing the number of unique and common proteins detected in a native pregnancy mucus plug (*21*) versus mucus cleaved from Endocervix Chips (N=2 human donors) using proteomic analysis. **C**) Bar graph showing differential abundance of undecorated, sialylated, sialofucosylated (SF), and sulfated O-glycans in mucus cleaved from Endocervix Chips versus Ectocervix Chips determined using nano LC-MS glycomic profiling. Structures are grouped based on glycan composition and retention time (Hex_HexNAc_NeuAc_Fuc| RT) (N=3-4 chips per condition; Mean ± SD, one-way ANOVA; E2= estradiol, PH= progesterone and estradiol, \**p*<0.05, \*\**p*<0.01, \*\*\**p*<0.001).

Although there is limited data on the mucin composition or structure of native human endocervix during pregnancy, a human mucus plug proteome has been published (*21*). To evaluate how closely the Endocervix Chip reproduces native mucus plug composition, we carried out unbiased proteomic analysis of mucus isolated from the apical channel of the PAH-treated Endocervix Chip and confirmed that it is a rich reservoir of proteins and peptides (**Fig. 2B**). Of the 3639 proteins found within enzymatically cleaved mucus from chips, 716 were also previously detected in native pregnancy mucus plugs (*21*). These shared proteins included mucins MUC1, MUC5B, MUC5AC and MUC16, AMPs (e.g., S100A9), whey acidic protein (WAP), four-disulfide core domain protein 2 (WFDC2), SAA1/2, elafin (PI3), lactotransferrin (LTF), lysozyme (LYZ) and secretory leukocyte protease inhibitor (SLPI). The published proteome of the native mucus plug contained 414 additional proteins that were not detectable in the Endocervix Chip plugs; however, many of these proteins are known to be produced by either immune cells (e.g. MMP-9, DEFA) or by other tissues within the female reproductive tract not present in our chips. These results show that the human Organ Chip approach can provide insights into the relative contributions of specific tissues and cell types to mucus plug formation and hence, host defenses to infection.

As glycans within the mucus are key modulators of host-microbiome interactions (*26*), we also carried out LC-MS/MS analysis, which revealed that both the epithelial cell type and hormonal milieu influence the mucus glycan profile in these cultures. Approximately 145 N-glycans isomers were identified as either sulfated, fucosylated, sialofucosylated or undecorated (**Fig. S4A).** The Endocervix Chips exhibited a higher proportion of sialylated structures and a marked decrease in sialofucosylated species, while fucosylated glycoforms were similar when compared to Ectocervix Chips. Importantly, exposure to the PAHs also significantly altered the N-glycan profile of the Endocervix chips, with a substantial reduction in the solely sialylated species being most notable. More pronounced differences in structure also were observed in O-glycan profiles (**Fig. 2C & fig. S4B**) as only the Ectocervix Chips contained certain sialofucosylated O-glycan structures while the Endocervix hips contained additional undecorated, branching core O-glycans (e.g., 4_4_0_0|25.3, 3_3_0_0|21.2, and 5_5_0_0|25.6) and a higher proportion of sialylated and sulfated O-glycans. Moreover, the relative abundances and ion counts of all undecorated O-glycans in Endocervix chips decreased upon hormonal treatment. Notably, the glycan 2_2_0_1|24.0 was significantly more abundant in the Endocervix Chip and its abundance was further increased with PAH treatment. Sulfation on sialylated structures, exemplified by glycan 2_2_0_1|29.6, also was observed exclusively in Endocervical Chips and similarly increased upon exposure to PAHs.

### Dysbiotic microbiome stimulates cervical inflammation and mucus plug breakdown

While preterm birth has been associated with the presence of a dysbiotic (non-optimal) microbiome (*3*, *27*), it is impossible to study the direct effects of the vagino-cervical microbiome on endocervix dysfunction during pregnancy. We therefore leveraged the human Endocervix Chips exposed to either β-estradiol (E2) or PAHs to mimic pregnancy and evaluated how they respond when co-cultured with a previously described healthy (optimal) vaginal microbiome composed of a consortium of patient-derived *Lactobacillus crispatus* strains (C6) versus a dysbiotic Bacterial Vaginosis Consortium 1 (BVC1) containing *Gardnerella vaginalis E2* and *E4*, *Atopobium vaginae* and *Prevotella bivia* (*28*).

Interestingly, the epithelial microenvironment of the Endocervix Chips chips appeared to more effectively suppress the growth of the dysbiotic BVC1 microbiome than that of the Ectocervix Chips. Quantification of bacterial load by digesting the epithelium with enzymes revealed that there were significantly lower levels of BVC1 colony forming units (CFUs) per chip at 96 hours of co-culture in the Endocervix Chips compared with the Ectocervix Chips (**Fig. 3A**). Moreover, there was even greater bacterial growth suppression when the Endocervix Chips were perfused with PAHs to mimic pregnancy, although the bacteria were not completely eliminated (**Fig. 3A**). These results are consistent with the known antimicrobial functions of the native endocervix and mucus plug (*6*) and the observation that the female upper reproductive tract has a much lower bacterial load than the vagina (*29*). In addition, these results underscore the benefits of using primary human cells from known anatomical locations within the female reproductive tract in research focused on host-microbiome interactions.

**Figure 3.**
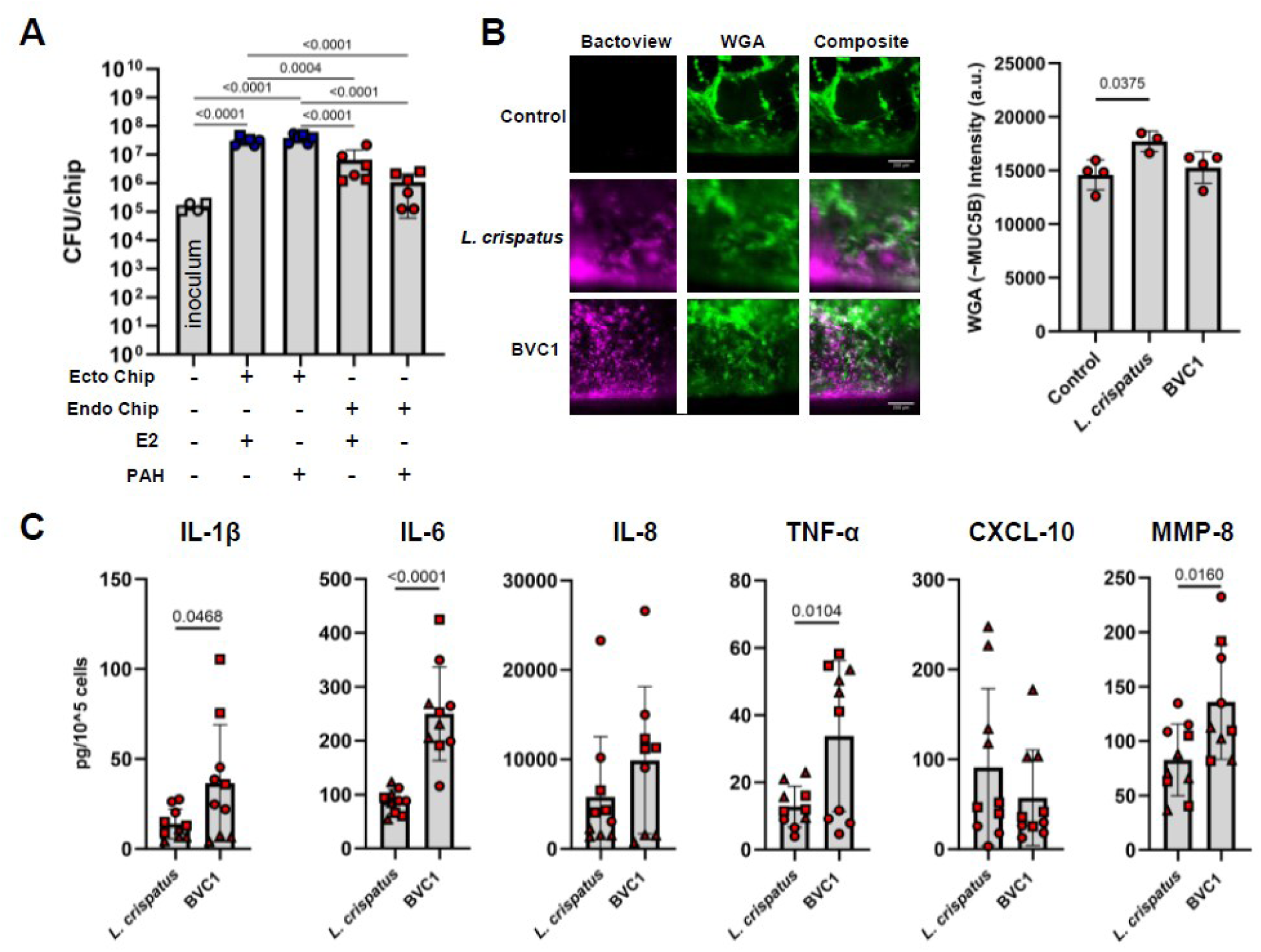
Cervix Chip-Vaginal Microbiome Interactions. **A)** Bar graph showing total colony forming units (CFUs)/Chip of BVC1 present in the inoculum and detected on Endocervix versus Ectocervix Chips 96 hours post-inoculation (N=6 chips per group, 2 independent experiments for each tissue type with each indicated by ○ or □ (Mean ± SD, log10 scale, one-way ANOVA). **B)** Immunofluorescence microscopic side-view images (left) of Endocervix Chips inoculated with either *L. crispatus* or BVC1 showing bacteria (purple, Bactoview) and mucus (green, WGA) (Bar, 200 μm) with bar graph at right depicting WGA lectin intensity (N=3-4 chips Mean ± SD, one-way ANOVA). **C)** Bar graphs showing cytokine (IL-1β, IL-6, IL-8, TNF-α, CXCL-10) and MMP-8 levels in apical outflows of Endocervix Chips 72 hours post-inoculation with the *L. crispatus* or BVC1 consortium (N=10 chips; 3 donors with results from each labeled as ○, □ or Δ; Mean ± SD, *t*-test).

As breakdown of the mucus plug(*30*) is a key feature of cervical dysfunction that can lead to preterm labor, we explored how a healthy versus dysbiotic microbiome influences mucus accumulation on-chip. The presence of the healthy *L. cripatus-*rich C6 consortium increased MUC5B levels, which filled the lumen of the epithelial channel to a much greater degree than observed in microbiome-free controls (**Fig. 3B**). In contrast, exposure to the dysbiotic BVC1 consortium resulted in less accumulation of mucus, which appeared as a loose network with bacteria appearing throughout (**Fig. 3B**). Finally, in addition to affecting the mucus plug-like structure on-chip, the presence of a dysbiotic BVC1 microbiome resulted in higher levels of secretion of the pro-inflammatory cytokines, IL-1β, IL-6 and TNF-α as well as the ECM-degrading MMP-8 enzyme compared to the healthy *L. crispatus* rich C6 consortium; however, the magnitude of the TNF-α inflammatory response varied between different donors (**Fig 3C**).

### Inflammatory cytokines induce cervical dysfunction and ECM breakdown

Recent studies combining metagenomic and cytokine analyses conducted as part of the integrative human microbiome project have shown that women with low levels of *L. crispatus* and elevated vaginal levels of pro-inflammatory cytokines IL-1β, IL-6, MIP-1β, and eotaxin are at increased risk to deliver preterm (*3*). Injected inflammatory cytokines also can trigger labor in rodents by activating uterine contractions (*17*); however, much less is known about the effects of inflammatory cytokines on the human cervix during pregnancy. We therefore perfused Endocervix Chips apically with inflammatory cytokines IL-1β (4 ng/ml), IL-6 (1 ng/ml), IL-8 (15 ng/ml), and TNF-α (0.2 ng/ml) at concentrations similar to those previously reported in vaginal secretions of women who delivered preterm (*27*), while continuously perfusing the basal channel with medium containing PAHs to mimic the systemic hormonal milieu of pregnancy. Exposure to these inflammatory cytokines upregulated epithelial production of MUC5AC in the Endocervix Chips as confirmed by staining with Ulex Europaeus Agglutinin I (UEA I) lectin(*31*) that binds to α-linked fucose residues and has been shown to co-localize with this mucin (*39*) (**Fig. 4A**). Increased expression of *MUC5AC* was confirmed by RT-qPCR, although there was no significant change in expression of *MUC5B* (**Fig. 4B**).

**Figure 4.**
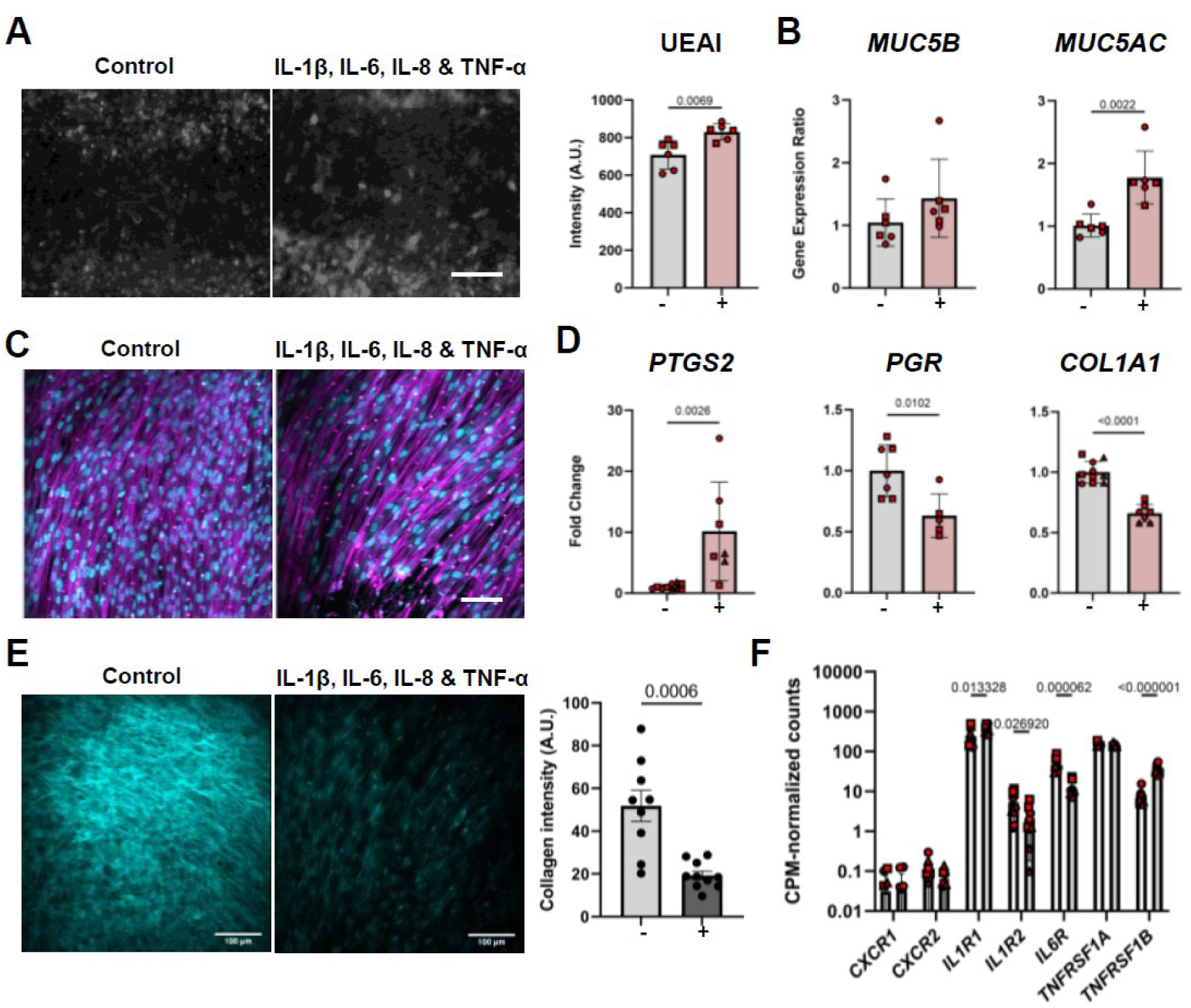
Effect of inflammatory cytokines in Endocervix Chips. **A)** Immunofluorescence microscopic top-view images of the epithelium within Endocervix Chips showing Ulex Europaeus Agglutinin I (UEA I)-stained mucus after 24 hours without (control) or with exposure to a cytokine mixture of IL-1β, IL-6, IL-8 & TNF-α (bar, 180 μm) and bar graph at the right showing quantification of fluorescence intensities (-, control; +, with cytokines; 6 non-overlapping fields of view; N=3 chips; Mean ± SD, *t*-test). **B)** Graph showing levels *MUC5B* and *MUC5AC* gene expression determined by qRT-PCR in Endocervix Chip epithelia 24 hours after perfusion with cytokine mixture shown in **A** (N=6 chips, 2 donors, 2 independent repeats, Mean ± SD, *t*-test). Data were normalized to housekeeping genes and control chips (ΔΔCt). **C)** Immunofluorescence microscopic images of the Endocervix Chip stroma stained with F-actin (magenta) and Hoechst (nuclei, cyan) (bar, 100 μm). **D)** Graph showing levels of *PTGS2*, *PGR*, and *COL1A1* gene expression in PAH-perfused Endocervix Chip stroma with (+) and without (-) cytokine treatment (N=6-10 chips, 2-3 donors with results from each donor labeled as ○, □ or Δ;). Data normalized to housekeeping genes and control chips (ΔΔCt, Mean ± SD, *t*-test). **E)** Representative second harmonic microscopic images (left) of fibrillar collagen within the stroma in the basal channel of control and cytokine-perfused Endocervix Chips (bar, 100 μm) with quantification shown in the bar graph at the right (-, control; +, with cytokines; 9-10 non-overlapping fields of view, N=3 chips, Mean ± SD, *t*-test). **F)** Gene expression levels expressed as count per million (CPM)-normalized counts of IL-8 (*CXCR1*, *CXCR2*), IL-1 (*IL1R1*, *IL1R2*), IL-6 (*IL6R*), and TNF-α (*TNFRSF1A*, *TNFRSF1B*) receptors in the Endocervix Chip epithelium (white bars) and stroma (light grey). (N=10 chips, 3 donors, Mean ± SD, Two-way ANOVA with multiple comparisons).

Interestingly, exposure to these preterm labor-associated inflammatory cytokines also altered the appearance of stromal cells in the basal channel, which became more aligned (**Fig. 4C**). Stromal expression of the prostaglandin producing enzyme *PTGS2* was also upregulated, while expression of both stromal collagen *COL1A1* and progesterone receptor (*PGR)* were reduced (**Fig. 4D**). These are important findings because prostaglandins are routinely administered clinically to induce cervical ripening which is associated with collagen dissolution and induction of labor in pregnant patients, while the *PGR* is critical for maintaining hormone responsiveness and a decrease in its expression is suggestive of functional progesterone withdrawal (*34*). Similar effects were observed at the protein level, with a significant loss of fibrillar collagen being observed within the stroma as demonstrated by second harmonic imaging (**Fig. 4E**).

We found that endocervical epithelial and stromal cells only expressed low levels of the IL-8 receptors *CXCR1* and *CXCR2*; however, stromal cells expressed high levels of *IL1R1*, *TNFRSF1A*, and *IL6R*, suggesting that they may respond to these inflammatory signals (**Fig. 4F**). Notably, compared to the epithelium, the stroma of the Endocervix Chip expressed slightly lower levels of *IL1R2*, a decoy receptor that dampens inflammation by competing with *IL1R1* for IL-1 binding. This reduced expression of *IL1R2* suggests that the stroma may be more sensitive to IL-1–mediated inflammation. Additionally, the stroma exhibited higher levels of *TNFRSF1B*, which promotes pro-survival signaling, in contrast to *TNFRSF1A*, whose signaling drives apoptosis. These findings indicate distinct inflammatory and survival signaling dynamics between stromal and epithelial compartments in the Endocervix Chip. The results are also consistent with inflammatory cytokines inducing pathological processes involved in cervical dysfunction in both the epithelium and stroma, including changes in the mucus plug and premature cervical softening and shortening, which are associated with preterm labor.

### Immune cells amplify effects on cervical dysfunction

Because cervical remodeling associated with preterm labor involves immune cells(*35*), we also carried out studies in which peripheral blood mononuclear cells (PBMCs) were introduced into the Endocervix Chip to further increase the fidelity of *in vivo* mimicry in this model of cervical dysfunction. When inflammatory cytokines were perfused through the epithelial channel, followed by introduction of PBMCs into the basal channel one day later, equal numbers of PBMCs adhered to the stroma in both control and inflamed chips (**Fig. 5A**). Within 24 hours, they actively migrated through the stromal ECM and pores of the ECM-coated membrane, and into the epithelium above (**fig. S5A,B**). Importantly, even in the absence of inflammatory cytokines, inclusion of PBMCs on-chip led to a small but significant increase in secretion of MMP-8 and Cathepsin S into the basal channel (**Fig. 5B**). This suggests that PBMCs may contribute to tissue remodeling and repair, consistent with their roles in other organs (*36*).

**Figure 5.**
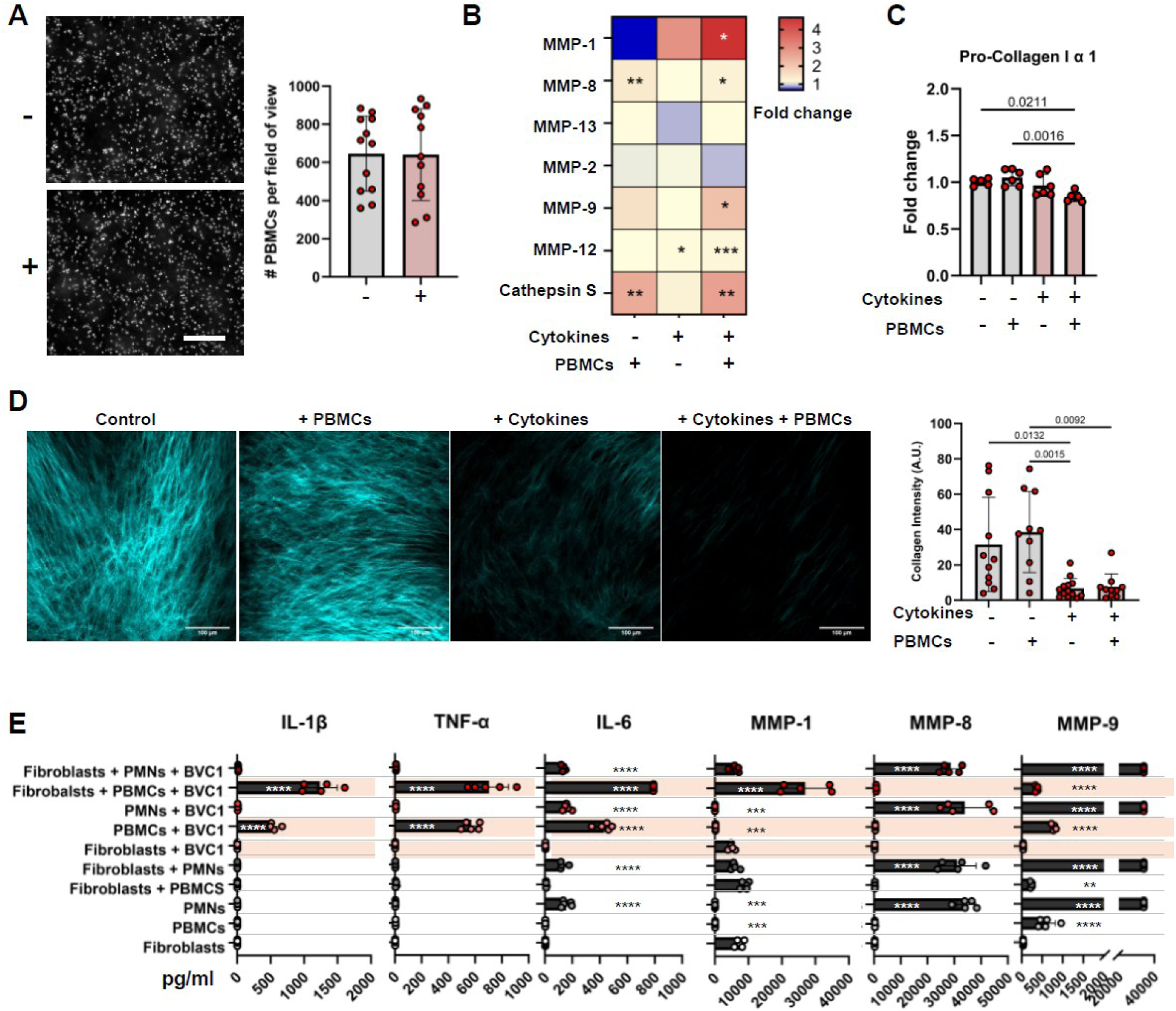
Effects of immune cells on stromal remodeling in Endocervix Chips. **A)** Immunofluorescence microscopic images (left) of the basal channel of an Endocervix Chip showing PBMCs (white) retained on the surface of the stroma when cultured in the absence (-) or presence (+) of IL-1β, IL-6, IL-8, and TNF-α for 5 hours (bar, 180μm, Mean ± SD, *t*-test) and graph at right showing quantification of the data (12 non-overlapping fields, N=4 chips, Mean ± SD, *t*-test). **B)** Heat map of matrix metalloproteinases (MMP-1, MMP-2, MMP-8, MMP-9, MMP-12, MMP-13) and Cathepsin S protein levels in basal channel outflows of Endocervix Chips cultured in the presence (+) or absence (-) of cytokines and/or PBMCS (data are normalized to controls without PBMCs; N=4 for both exposure to cytokines for 48 hours and PBMCs for 24 hours; one-way ANOVA, median values, significance levels relative to controls without PBMCs, \**p*<0.05, \*\**p*<0.01, \*\*\**p*<0.001). **C)** Bar graph showing pro-collagen 1α1 protein concentration fold change compared to control chips in basal outflows of Endocervix Chips cultured in the presence (+) or absence (-) of cytokines and/or PBMCS (N=5-6 chips, two independent repeats, Mean ± SD, one-way ANOVA). **D**) Representative second harmonic microscopic images (left) of fibrillar collagen within the stroma in the basal channel of Endocervix Chips cultured in the absence (control) or presence of PBMCs, cytokines, or both (bar, 100 μm) and bar graph at the right showing quantification of the results (N=3 chips,10-13 independent fields of view, Mean ± SD, one-way ANOVA). **E**) Bar graphs showing the concentrations of inflammatory cytokines (IL-1β, TNF-α, IL-6), collagenases (MMP-1, MMP-8) and gelatinase (MMP-9) measured in medium of conventional 2D cultures of fibroblasts, PBMC, or PMNs alone or in combination when co-cultured with or without the BVC1 consortium for 24 hours (N=5 wells, Mean± SD, one-way ANOVA, multiple comparisons, \**p*<0.05, \*\**p*<0.01, \*\*\**p*<0.001, \*\*\*\**p*<0.0001).

Moreover, when both PBMCs and inflammatory cytokines were present, the chips exhibited significantly elevated secretion of multiple ECM-degrading enzymes, including collagenases (MMP-1, MMP-8), gelatinases (MMP-9), elastases (MMP-12), and Cathepsin S, compared to unstimulated controls (**Fig. 5B, fig. S5C**). Incorporation of PBMCs into inflammatory cytokine-perfused chips also significantly lowered pro-collagen 1α1 levels compared to chips that were not exposed to inflammatory cytokines (**Fig. 5C**). While PBMCs alone did not lead to fibrillar collagen loss in the Endocervix Chip stroma, exposure to pro-inflammatory cytokines dramatically reduced fibrillar collagen signal, with PBMCs having no detectable protective or exacerbating effect (**Fig. 5D**).

Given that these findings contrast with prior studies suggesting macrophages and neutrophils are the primary effectors of cervical collagen degradation (*37*), we further investigated interactions between the endocervical stromal cells, PBMCs, polymorphonuclear cells (PMNs), and the non-optimal microbiome consortium BVC1 in a 96-well plate assay. Notably, PBMCs did not directly secrete MMPs in response to BVC1 but instead triggered a strong inflammatory response by the stromal cells, characterized by increased production of IL-1β, IL-6, and TNF-α (**Fig. 5E**). In contrast, PMNs appeared to secrete high levels of MMP-8 and MMP-9, although PMNs rapidly degranulate *in vitro* and MMP-9 levels were detected at the maximum limit of our assays; thus differences caused by exposure to BVC1 may have been masked in this assay.

Strikingly, only a tri-culture of PBMCs, stromal cells, and BVC1 led to a significant increase in MMP-1 secretion compared to stromal cells alone (**Fig. 5E**). This suggests that while PBMCs contribute to cervical dysfunction in the presence of microbial insult, their role may be primarily through triggering a stromal cell state change via inflammatory cytokine signaling, rather than through direct collagen degradation by the gelatinase MMP-9, as previously suggested(*38*). While PMNs secreted high levels of the collagenase MMP-8 in line with previous reports, they appeared to be acting directly, rather than via modulation of stromal cell activity in our assay.

### IL-1 is the critical inducer of changes in the endocervical stroma

Importantly, when we tested each cytokine (IL-1β, IL-6, or IL-8) individually, we discovered that IL-1β was solely responsible for the effects we observed. Only IL-1β significantly downregulated *COL1A1, COL3A1 and PGR* expression and upregulated *PTGS2* in the stromal cells when cultured in conventional planar culture dishes in the presence of either E2 or PAHs (**Fig. 6A**). When we analyzed effects on general MMP activity under similar E2 or PAH conditions, we found again that only IL-1β induced ECM-degrading enzyme activity (**Fig. 6B)**. Interestingly, PAH conditions resulted in overall lower levels of MMP activity in the presence of IL-1β, consistent with previous findings that progesterone decreases MMP stimulation by inflammatory cytokines in other cell types(*37*).

**Figure 6.**
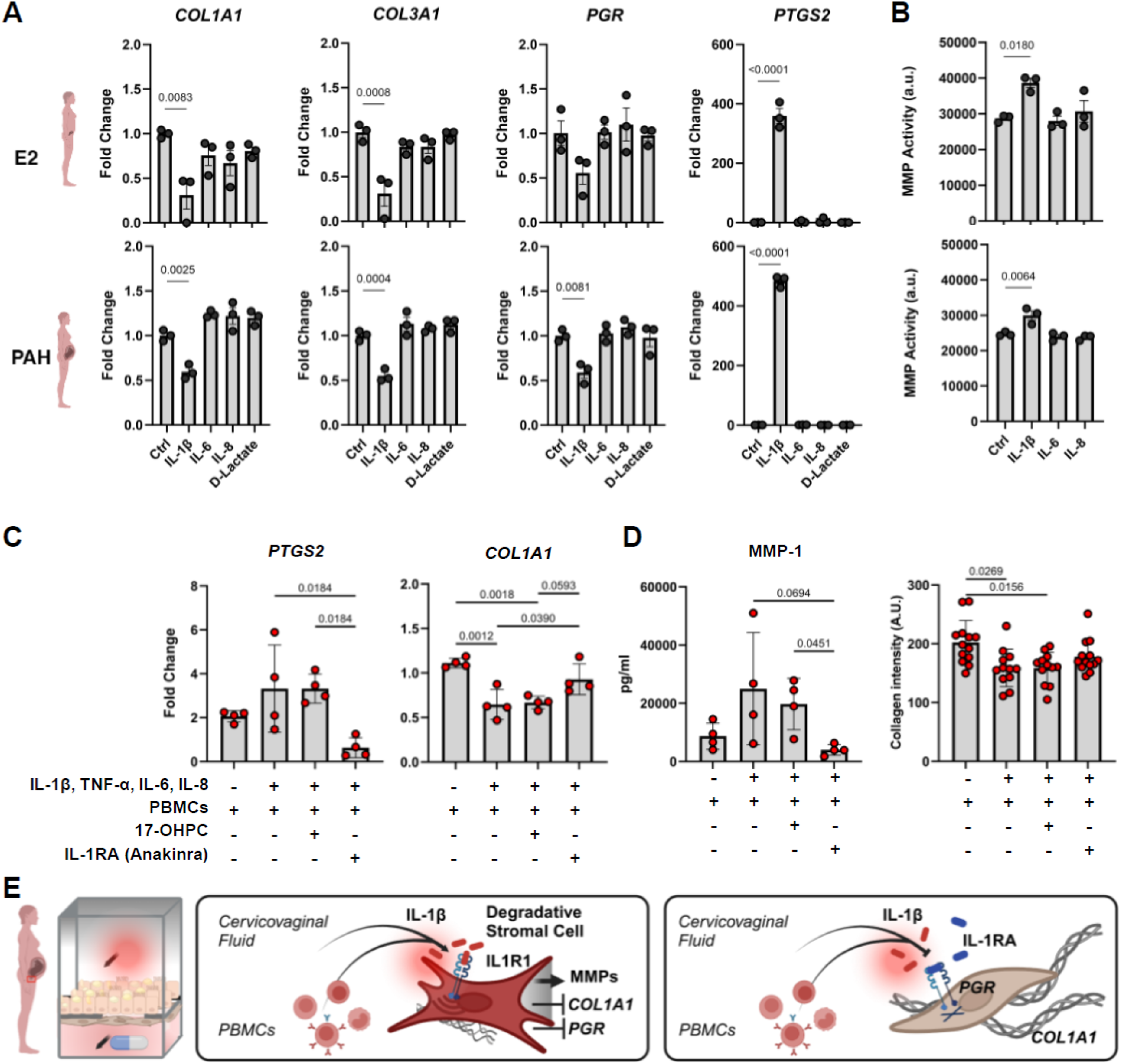
Drug testing in Endocervix Chips. **A**) Quantification of fold changes in expression of collagen (*COL1A1*, *COL3A1*), progesterone receptor (*PGR*), and prostaglandin synthase 2 (*PTGS2*) gene expression in endocervical fibroblasts cultured in culture medium with E2 or PAH in the absence (ctrl =control) or presence of IL-1β, IL-6, IL-8 or DL-Lactate for 24 hours (N=3 wells; Mean ± SD; ΔΔCt relative to housekeeping genes and control chips, one-way ANOVA, multiple comparisons). **B**) Bar graphs depicting total MMP activity in supernatants of endocervical fibroblast cultured for 24 hours in medium containing E2 (top) or PAH (bottom) in the absence (ctrl = control) or presence of individual inflammatory cytokines (IL-1β, IL-6, or IL-8) (N=3 wells, Mean ± SD, one-way ANOVA). **C**) Graphs showing stromal expression levels of preterm labor-associated genes (*PTGS2*, *COL1A1*) in Endocervix Chips cultured with PBMCs in the absence (-) or presence (+) of inflammatory cytokines (IL-1β, IL-6, IL-8, and TNF-α), 17-OHPC, or IL-1RA (Anakinra) for 72 hours. Expression is normalized to housekeeping genes and control chips without PBMCs (ΔΔCt) (N=4 chips, Mean ± SD, one-way ANOVA). **D**) Graphs showing MMP1 protein levels in the basal outflow measured 48 hours after cytokine exposure by Luminex (N=4 chips, Mean ± SD, one-way ANOVA) (left) and fibrillar collagen intensity levels measured 72 hours after exposure to cytokines by second harmonic imaging (N=3-4 chips per group. 12-15 non-overlapping fields, Mean ± SD, one-way ANOVA) when Endocervix Chips were cultured under similar conditions as shown in **C**. **E**) Schematic representation of inflammation-induced changes in endocervical stromal cells and the therapeutic reversal of this phenotype via administration of an IL-1 receptor antagonist (IL-1RA). Chip diagram (left) depicts inflammatory cytokines (red circle) being perfused through the apical channel to mimic local inflammation and drug perfused basally to mimic systemic delivery. Middle schematic shows that IL-1 (red ovals) switches the state of stromal cells to degradative stromal cell phenotype marked by an increased expression of MMPs and reduced expression of *COL1A1* and *PGR*. Blocking IL-1R1-mediated signaling (right schematic) via IL-1RA (blue ovals) prevents cervical fibroblast activation and preserves collagen (triple helix). Created with BioRender.

### Assessing the efficacy of potential therapeutics for preventing cervical dysfunction

Having established that preterm labor-associated inflammatory cytokines result in ECM dissolution within the stroma of the Endocervix Chip, we next explored whether this could be prevented by administering anti-inflammatory medications. Our studies showed that cytokines induce expression of *PTGS2*, which encodes the cyclooxygenase-2 (COX-2) enzyme that produces prostaglandins which are also used clinically to promote cervical ripening and accelerate labor. Thus, we first evaluated whether the COX-2 inhibitor drug indomethacin that is used clinically to treat preterm labor could improve cervical integrity on-chip. Interestingly, we found that indomethacin prevented the increase in *PTGS2* expression in PAH-perfused Endocervix Chips exposed to inflammatory cytokines, but it did not prevent the cytokine-induced decrease in *COL1A1* and *PGR* expression (**fig. S6A)**.

We next tested hydroxyprogesterone caproate, which remains the standard of care for preventing preterm labor in many countries even though the FDA recently rescinded its approval for clinical use for this application because of failure to convincingly demonstrate efficacy (*39*). In line with the FDA’s findings, we found that hydroxyprogesterone caproate (17-OHPC) did not suppress the effects of inflammatory cytokines on stromal cells that are associated with promotion of cervical dysfunction (i.e., increased *PTGS2*, decreased *COL1A1*, and increased MMP-1) when measured on-chip (**Fig 6C,D**). In contrast, when we tested the recombinant human IL-1 receptor antagonist (IL-1RA), anakinra, we found that it significantly decreased the expression of *PTGS2* and increased the expression of *COL1A1* compared to controls (**Fig. 6C)**. Similarly, only anakinra reduced the release of MMP1 into the basal medium and prevented the reduction of fibrillar collagen in response to cytokine exposure (**Fig. 6D)**. Taken together, these results support our finding that IL-1β is the primary cytokine responsible for driving cervical dysfunction in this model and that IL-1R1 inhibitor drugs, such as anakinra, may be useful for preventing preterm labor in humans (**Fig. 6E**).

## DISCUSSION

Cervix dysfunction is a recognized risk factor for preterm labor. Yet, molecular processes leading to cervix dysfunction and how to prevent it remain poorly understood due to the lack of suitable experimental approaches. In this article, we described microfluidic Organ Chip models of human endocervix that contain both mucus plug-producing epithelial cells and ECM-producing stromal fibroblasts as well as circulating immune cells, which recapitulate key organ-level functions relevant to cervical dysfunction. Transcriptomic analysis confirmed that the epithelium within the Endocervix Chips express relevant structural and functional genes including the cytokeratin *KRT7* and mucins *MUC5B* and *MUC5AC,* while the stromal cells deposited a rich fibrillar ECM containing *COL1A1* and *COL3A1*, key constituents of the endocervical ECM *in vivo* (*40*). The ability of the endocervical stroma to self-assemble its own ECM on-chip is a major advantage of our model, as stromal remodeling is mediated by changes in ECM composition and crosslinking (*41*), which can differ significantly in different tissue types and commercial ECM formulations. Importantly, untargeted proteomic analysis of lumenal secretions in PAH-perfused Endocervix Chips also confirmed that a mucus plug was produced, which has a composition that is highly similar to that of the native human mucus plug (*21*). This level of similarity has been a long-standing challenge to achieve *in vitro*, and it includes production of several mucins (MUC1, MUC5B, MUC5AC, MUC16) and AMPs (WFDC2, SAA1, S100A9, PI3, LTF, LYZ, SLPI) on-chip.

The vaginal microbiome is abundant in the lower reproductive tract while the upper reproductive tract is almost sterile (*29*), with the cervical mucus plug playing a key role in protecting the fetus from ascending infections(*31*). Our study suggests that this could, at least in part, be due to the production of distinct AMPs, glycans, and nutrients expressed and secreted by the functionally distinct endo– and ecto-cervical epithelia. Interestingly, we found that the dysbiotic BVC1 vaginal microbiome grew significantly more rapidly in Ectocervix Chips compared with Endocervix Chips, especially when chips were perfused with PAHs. This may relate to differences in the defense mechanisms of these two different anatomical regions of the cervix. For example, WAP four-disulfide core domain protein 2 (*WFDC2)* that was expressed at higher levels in Endocervix Chip compared to the Ectocervix Chip was recently shown to potently reduce the viability of *Enterococcus faecalis*, *Staphylococcus aureus*, *Escherichia coli* and *Pseudomonas aeruginosa* (*42*). In addition, mucins MUC5B and MUC5AC that were present at much higher levels in the Endocervix Chip are known to alter the composition of oral bacterial consortia (*43*) and reduce the virulence of pathogens. Similarly, the *N*– and *O*-glycan profiles of the Ecto– and Endocervix chips were distinct, which could influence microbial growth as well. Further understanding of the antimicrobial networks that keep the cervicovaginal microbiome in check will be important as we previously showed that transferring outflows from Cervix Chips lined by a mixture of endo– and ecto-cervical epithelia to Vagina Chips significantly reduced the levels of the BVC1 consortium (*28*). The current primary Endocervix Chip model also revealed that despite the protective functions of its secretions and the mucus plug it forms, the presence of a dysbiotic microbiome still resulted in an elevated inflammatory response compared to the healthy *L. crispatus* consortium. This aligns well with the body of literature and clinical reports documenting adverse cervico-vaginal symptoms associated with a non-optimal microbiome (*3*, *27*).

Several clinical studies suggest a link between elevated cervico-vaginal cytokines related to bacterial dysbiosis and preterm labor (*3*, *27*). However, studying preterm labor has been notoriously challenging due to the limitations of existing models and ethical concerns. For example, a standard rodent preterm labor workflow involves injection of either lipopolysaccharide endotoxin (LPS) or inflammatory cytokines into the uterine horn or intraperitoneally, which rapidly triggers contractions and preterm labor(*44*). However, parts of the labor cascade, such as the premature rupture of membranes or cervical shortening often precede myometrial activation in human preterm labor. Here, we present direct evidence that exposure of endocervical epithelium to inflammatory cytokines at concentrations observed in the cervicovaginal fluids of women who deliver preterm within the organ-relevant environment of the Endocervix Chip leads to significant pathological alterations of the endocervix. First, inflammatory cytokines alter mucus plug composition, notably by increasing MUC5AC levels.

These results are consistent with the effect of inflammation on altered mucus production in other mucosal tissues (*45*). Exposure to inflammatory cytokines also induces degradation of the collagenous ECM as confirmed by second harmonic imaging. Importantly, a similar decrease in collagen content was observed by others using whole human cervix imaging throughout pregnancy in women who delivered preterm (*46*). In contrast, in our *in vitro* model we were able to gain insight into the underlying mechanism by showing that the reduction in total collagen content is due to both downregulated *COL1A1* and *COL3A1* expression and enhanced MMP secretion by endocervical stromal cells.

It is also important to note that IL-1β downregulated progesterone receptor levels in the stroma in our studies, which is indicative of functional progesterone withdrawal. Interestingly, cervical fibroblasts from women with refractory cervical insufficiency have consistently impaired responses to progesterone (*47*). These results align with prior reports conducted with cervical smooth muscle cells in conventional planar cultures (*48*), however, in addition they demonstrate the functional impact of these changes on clinically relevant readouts. Another novel finding is that the dissolution of endocervical ECM we observed in response to inflammatory cytokines did not require the presence of immune cells. While our results are consistent with the notion that immune cells contribute to cervical remodeling (*38*), they suggest that cervical dysfunction may be due to direct stromal activation by immune cell-secreted inflammatory cytokines resulting in a stromal cell state shift to a degradative phenotype.

Having established a reliable model of human endocervix dysfunction, we explored how this model can be utilized to screen potential therapeutic strategies. This issue is especially pressing due to the lack of effective treatments to prevent preterm labor, which culminated in the FDA’s 2023 withdrawal of approval for the progestin-based drug Makena (hydroxyprogesterone caproate or 17-OHPC). In our model, we also found that 17-OHPC had no protective activity and the NSAID indomethacin, while reducing inflammation, failed to prevent tissue damage. In contrast, our results suggest that the cytokine-specific anti-inflammatory drug anakinra, which targets IL-1 signaling, may be more effective than both NSAIDs and progestins in preserving cervical integrity and preventing preterm labor by directly modulating cytokine-driven ECM remodeling. Further investigation into the safety and efficacy of IL-1R antagonists in clinical settings is warranted, especially since these drugs also prevent myometrial contractions in a human-specific model (*18*). Taken together, these results suggest that the human Endocervix Chip model offers a promising platform for testing therapeutic interventions (*49*), which could potentially accelerate the discovery of effective treatments (*49*) for preterm labor. More broadly, these results also provide a potential mechanistic link between studies showing a close association between elevated vaginal immunoproteome and spontaneous preterm birth (*50*).

Despite these advances, our study has limitations, including mimicking pregnancy using a synthetic combination of female hormones. We also used medroxyprogesterone acetate instead of native progesterone and the dose of this hormone used at baseline may mask some effects of progestin-based drugs with their known anti-inflammatory properties (*22*, *51*). We also tested a low TNF-α concentration as clinical studies do not report this cytokine to be elevated in the context of microbiome-related preterm labor (*3*). It also would be interesting to incorporate tissue resident immune cells, a broader range of cytokines (*3*) and other preterm labor triggers (e.g. LPS, *Escherichia coli),* as well as potentially plasma from pregnant donors in future studies to fully capture the complexity of preterm labor and its different subtypes.

In conclusion, we have developed a primary Endocervix and Ectocervix Chip models that enable new insights into how these different anatomical regions of the human cervix interact with healthy and dysbiotic microbiome, respond to inflammatory cues, and influence cervical dysfunction in a pregnancy-like environment. This platform holds great potential for identifying new treatment strategies aimed at preventing preterm birth, shifting the focus from managing preterm delivery to early detection and prevention.

## MATERIALS AND METHODS

### Study Design

The goal of the study was to develop a tool to study human cervix dysfunction related to preterm labor. The primary goals of this study were confirming that the Cervix organ chip model recapitulates the key features of the endocervix: mucus production and collagen deposition. Identity of the cells was confirmed using immunohistochemistry and RNAseq and involved collaboration with experienced pathologists. Multiple complementary assays were selected to confirm this and at least two different human donors were used to confirm the methods described in this study result in the relevant phenotype. Cervix dysfunction was defined as a loss of collagenous ECM and was confirmed using multiple distinct assays in chips derived from at least two distinct donors. Contributions of IL-1 beta to cervix dysfunction were studied both using the Endocervix chips and also using conventional cell-culture dishes to ensure the results are reproducible across different experimental systems.

### Effect of cytokines and Indomethacin in Endocervix chips

Endocervix chips were differentiated under the PAH hormonal condition (**Supplementary Table 1**). To mimic inflammation in adjacent tissues, the endocervix chips were apically perfused with a mixture of IL-1β (4ng/ml) (Stem Cell Technologies), IL-6 (1 ng/ml) (Stem Cell Technologies), IL-8 (15 ng/ml) (Stem Cell Technologies) and TNF-α (0.2 ng/ml) (Abcam, AB52126) in HBSS++ supplemented with Primocin for 4 hours daily followed by 20 hours of stasis for up to 72 hours. Some chips were additionally pre-treated for 80 minutes and then constantly perfused with 15 µM Indomethacin (Thermo, J3255.03) in the basal channel.

### Effect of cytokines and Anakinra in Endocervix Chips

The Endocervix chips differentiated under PAH conditions were perfused basally with either 10μg/ml Anakinra (Raleukin; FisherScientific; 50-202-9648), 1μM Hydroxyprogesterone Caproate (VWR, 103544-986) or left untreated all in the presence of PAH hormones for 24hours. At 24 hours, some groups were additionally perfused apically with a mixture of IL-1β (4ng/ml) (Stem Cell Technologies), IL-6 (1 ng/ml) (Stem Cell Technologies), IL-8 (15 ng/ml) (Stem Cell Technologies) and TNF-α (0.2 ng/ml) (Abcam AB52126) in HBSS++ for 4 hours daily at 40μl/h while continuing the basal treatment. 24 hours after the initial cytokine exposure, the chips were incubated with PBMCs that were pre-stained with CellTracker^TM^ Deep Red (ThermoFisher Scientific, C34565). 25μl of PBMCs resuspended in basal medium with relevant treatment at the concentration 50×10^6^/ml were injected into the basal channel and allowed to adhere for 2 hours to facilitate adhesion, perfused at 100 μl/h for 20 minutes to remove unbound PBMCs and thereafter exposed to 0/40 μl overnight flow. After 48 hours post cytokine exposure, basal outflows were collected for analysis. Stroma was visualized at 72 hours post first cytokine exposure using second harmonic imaging and the chips were lysed using Buffer RLT Plus for gene expression analysis (Qiagen, 1053393).

### Data analysis, Statistics and Visualizations Software

Data analysis, statistics and visualization was performed using the GraphPad Prism 10.1.0 software. Images were processed using ImageJ (FIJI, 2.1.0/1.53c, Java 1.8.0_172) using the Presentation plugin (Imperial College London) and the Imaris software. Omics data analysis is described in Supplementary materials. Diagrams were created using Biorender (https://app.biorender.com/).

### Statistical Analysis

Each chip was used as a biological replicate and experiments were typically repeated multiple times with different donors. As part of routinely implemented quality control, fluid flow through chips was monitored and chips and sampling time points that had perturbed flow (e.g. mixing of fluid between channels) were excluded from the final analysis. Either a Student’s *t*-test, one-way ANOVA or two-way ANOVA were performed to determine statistical significance and are further described in individual figure legends. Error bars are plotted as standard deviation (SD) and values p<0.05 were considered as significant.

### List of Supplementary Materials

Materials and Methods

Fig S1 to S7

Tables S1 to S5 for multiple supplementary tables

References (*01–05*)

## Acknowledgments

We thank Marina Feigenson, Ph.D., Yuncheng Man, Ph.D., Alican Ozkan, Ph.D, Pranav Prabhala and Ryan Posey for donating PBMCs for this project and Hassan Rhbiny for initial support. We also thank the VMRC consortium, especially Jacques Ravel, Ph.D, and Seth Rakoff-Nahoum, Ph.D., for providing the *L. crispatus* and *G. vaginalis* strains used in this study.

## Funding

Gates Foundation (INV-035977)

Wyss Institute for Biologically Inspired Engineering at Harvard University

## Author contributions

Conceptualization: AS, DEI

Methodology and Investigation: AS, KC, DBCH, AG, JF, ZI, OG, YB, SCH, RP, JC, JH, BB, AJ, CL

Visualization: AS, KC NB, SCH, TF, JF

Funding acquisition: AS, AJ, GG, DEI

Project administration: AJ

Supervision: DEI, CBL

Writing – original draft: AS, DEI;

Writing – review & editing: All authors

## Data and materials availability

All data are available in the main text or the supplementary materials. Proteomics data was deposited in the MASSIVE public dataset (MSV000097666). RNAseq data was deposited to the Gene Expression Omnibus (GE) database with the accession number GSE295361.

## SUPPLEMENTARY INFORMATION

### List of Supplementary Materials

Materials and Methods

Fig S1 to S7

Tables S1 to S5 for multiple supplementary tables

References (*01–05*)

## MATERIALS AND METHODS

### Primary Epithelial Cell Isolation

Endocervix and Ectocervix specimens were anonymously collected from patients undergoing hysterectomy for benign conditions (uterine fibroids, endometriosis, menorrhagia) at the Massachusetts General Hospital under IRB-approved protocol #2015P001859 or obtained from organ donors through the International Institute for the Advancement of Medicine (IIAM). Following a wash step in Hanks’ Balanced Salt Solution (HBSS++; Thermo Fisher; 14025092) or Phosphate Buffered Saline (PBS-/-; Gibco, 14190144) with Primocin (InvivoGen, ant-pm-1), the surgical explanted endocervical tissues were incubated in a solution containing 50% 5U/ml Dispase (STEMcell technologies, 07913) and 50% advanced DMEM/F12 basal medium (ThermoFisher Scientific, 12634010) with Primocin overnight at 4°C (alternatively the tissue may be incubated at 37 °C for several hours) to enzymatically separate the epithelium from the stroma. The epithelium was then mechanically scraped away from the stroma using a Cell scraper (FALCON, 353085) and the tissue was rinsed with PBS –/-between scrapings in order to collect as many epithelial cells as possible. The epithelial cells were collected in a falcon tube, spun down at 1500 RPM for 5 minutes, and the supernatant was discarded. Subsequently, to separate the epithelial sheets into single cells, the epithelial cells were treated with TrypLE™ Express Enzyme (1X) (ThermoFisher Scientific, 12604013) for 15 minutes at 37°C. Finally, the isolated endocervical epithelial cells were spun down at 1500 RPM for 5 minutes, supernatant was discarded, and cells were plated on T-75 or T-150 flasks depending on cell yield. Primary Cervical Epithelial Cells (ATCC® PCS-480-011™) from a batch that we confirmed was only KRT7 positive were used for the drug evaluation experiment. All endocervical epithelial cells were expanded in an expansion medium containing 50% conditioned WRN medium, Advanced DMEM/F12 and other factors listed in (**Supplementary Table 2**) that was based on the protocol described by Chumduri and colleagues(*13*) prior to cryopreservation. Ectocervical epithelial cells were isolated from ectocervical donor tissue following the same protocol described above and were grown in the DermaCult™ Keratinocyte Expansion Medium supplemented with hydrocortisone (STEMCELL Technologies, 100-0500) on T-75 or T-150 flasks prior to biobanking.

### Primary stromal cell isolation

For both the endocervix and ectocervix, following epithelial isolation, the stroma was minced into small pieces using a surgical scalpel/dissection scissors and digested in a digestion buffer containing 0.1% (w/v) collagenase type I (Worthington, LS004196) and 0.25% (w/v) collagenase type II (Worthington, LS004176) in Advanced DMEM/F12 basal medium with Primocin overnight at 37°C (alternatively the tissue may be incubated at 37 °C for several hours). The resulting stromal cell suspension was then centrifuged at 1500 RPM, and supernatant was discarded. The isolated stromal cells were grown in DMEM(1) + GlutaMAX^TM^-I, 4.5g/L D-Glucose (gibco, 10569-010) supplemented with 10% FBS and Primocin on T-75 or T-150 flasks for expansion prior to biobanking.

### Endocervix Chip Culture

Commercially available 2-channel Organ Chips devices (S1 Chip; Emulate Inc.) were first activated and washed (according to chip manufacturer protocol) then incubated with 500 μg/ml Collagen IV in 0.25% acetic acid (Sigma, C7521) in the top channel and a mixture of 200 ug/ml Collagen I (Corning, 354236) and 30 ug/ml Fibronectin (Corning, 354008) diluted in phosphate buffered saline ((PBS –/-) Gibco, 14190144) in the bottom channel overnight at 37°C to coat the porous membrane separating the two channels. The chips were then washed with DMEM + GlutaMAX^TM^-I (Gibco, 10569-010) supplemented with 10% FBS (fibroblast medium) and the basal side of the membrane was seeded with endocervical stromal cells by introducing 40μl of a cell suspension (6 x 10^5^ cells/ml). To ensure stromal cells adhered to the bottom of the porous membrane, the chips were inverted and incubated at 37°C for 3 hours in a humidified 5% CO_2_ incubator before being sealed with medium containing pipette-tips. The next day, the basal channel medium was replaced with low serum fibroblast medium (DMEM(1) + GlutaMAX^TM^-I, gibco, 10569-010, supplemented with ATCC fibroblast growth kit-low serum (ATCC, PCS-201-041). The apical channel was then seeded with primary endocervical epithelial cells (2 x 10^6^ cells/ml), in a mixture of Cervical WRN-Medium (25%) (**Supplementary Table 2,3**) and DermaCult™ Keratinocyte Expansion Medium (StemCell Technologies, # 100-0500) supplemented with Hydrocortisone Stock Solution (StemCell Technologies, 07925) and Primocin®. The chips were then maintained under static conditions for 3 days with the respective expansion medium in each channel being refreshed daily (**Supplementary table 3**). The chips were then connected to the ZOE instrument following manufacturer instructions (Emulate) and exposed to intermittent flow apically (40µl/hour) for 4 hours followed by 20 hours of stasis (0µl/hour)) and constant flow basally (40µl/hour) for a total of 48 hours to further expand the cell populations. To induce differentiation, the apical epithelial expansion medium was replaced with Hanks Balanced Salt Solution with calcium (HBSS++, (Thermofisher, 14025092)) supplemented with Primocin® (InvivoGen, ant-pm-1) and the basal stromal expansion medium was replaced with Advanced DMEM/F-12 medium (ThermoFisher Scientific, 12634010) supplemented with ReproLife^TM^ LifeFactors Kit (LifeLine, LS-1097) along with HLL supplement ( HSA 500 μg/mL, linoleic acid 0.6 mM, lecithin 0.6 μg/mL; ATTC), ascorbic acid (50 μg/mL; ATCC, PCS-201-040), and 5 nM β-Estradiol-Water Soluble (Millipore Sigma, E4389). To mimic pregnancy, following two days of basal perfusion with E2, the Pregnancy Associated Hormones-(PAH) chips were perfused basally with 45.884 nM β-Estradiol, 1 μM medroxyprogesterone acetate (Viovision, B2192-100), 2.5μg/ml Human Placental Lactogen (CSH1) (Sino Biologicals, 11596-H08H), 200 ng/ml Human Chorionic Gonadotrophin (HCG) (Abcam, 51782), and 140 ng/ml human recombinant prolactin (PeproTech®, 100-07-100ug). In the glycomics analysis, the PAH mixture consisted of the two key pregnancy hormones 0.1 nM β-Estradiol and 50 nM Progesterone P4 (Sigma, P7556). (**Supplementary Table 1**).

### Ectocervix Chip Culture

Primary ectocervical cells were expanded and differentiated on the Emulate Chips using similar approach to the Endocervix Chips, but using media known to support optimal Ectocervix phenotype. In particular, during the expansion stage the ectocervix epithelia (in the apical channel) were expanded in DermaCult™ Keratinocyte Expansion Medium supplemented with hydrocortisone (STEMCELL Technologies, 100-0500) and the ectocervix stroma (basal channel) was expanded in low serum fibroblast medium ((DMEM(1) + GlutaMAX^TM^-I,gibco, 10569-010), supplemented with ATCC fibroblast growth kit-low serum (ATCC, PCS-201-041)). To induce differentiation, the basal channel was perfused with the differentiation medium previously published by our group(*52*) supplemented with relevant hormones, while the apical channel was perfused with HBSS +/+.

### Effect of cytokines on primary endocervical fibroblasts in conventional cultures

Primary endocervical fibroblasts were treated with IL-1β (1 ng/ml) (Stem cell Technologies), IL-6 (10 ng/ml) (Stem Cell Technologies), IL-8 (50 ng/ml) (Stem Cell Technologies), Sodium DL-Lactate (5mM) (Santacruz, sc-301818B) additionally supplemented with either E2 alone or with the PAH mixture. Primary endocervical fibroblasts were seeded in a 12 well plate at 10,000 cells per cm^2^ in DMEM without phenol red (DMEM, high glucose, HEPES, no phenol red, Gibco, 21-063-029) supplemented with Fibroblast Growth Kit-Low serum (ATCC PCS-201-041,). On day 3, the medium was supplemented with E2. On day 4, the culture was supplemented with either E2 or PAH hormones in a medium without serum and without phenol red with the kit Fibroblast Growth Kit-Serum-free PCS-201-040™ from which the TGF-β supplement was removed. On day 6, the fibroblasts were treated with the relevant cytokines and the conditioned medium was collected for MMP activity analysis. Finally, cytokine treated primary endocervical fibroblasts were harvested using the RLT buffer (Qiagen, 79216) for gene expression analysis on day 7.

### MMP activity assay

MMP activity was assessed using a kinetic, MMP-sensitive fluorescent substrate cleaving assay. More specifically, supernatant was collected from primary endocervical stromal cells cultured in conventional well plates treated with inflammatory cytokines after 24 hours and MMP activity was evaluated using the SensoLyte® 520 Generic MMP Activity Fluorimetric Kit (Anaspec, AS-71158) based on manufacturer’s instructions. The MMPs were chemically activated as per manufacturers’ instructions and the activity was read on a plate reader. Readings obtained at 1 hour of the kinetic reaction were used for analysis.

### Co-cultures with microbiome on-chip

The dysbiotic consortium in this study termed Bacterial Vaginosis Consortium 1 (BVC1) consisted of *Gardnerella vaginalis E2* (C0011E2), *Gardnerella vaginalis E4* (C0011E4), *Prevotella bivia BHK8* (0795_578_1_1_BHK8), and *Atopobium vaginae* (0795_578_1_BHK4). The *Lactobacillus crispatus* consortium C6 consisted of three different strains (C0175A1, C0124A1, C0059A1) and was prepared from individual frozen aliquots as previously published(*52*).

Differentiated chips were infected apically with 37µl of inoculum resuspended in custom HBSS (LB/+G) as previously published(*52*) corresponding to approximately 10^5 colony forming units (CFUs) per chip and allowed to engraft under static conditions for 20 hours following which the chips were perfused with HBSS (LB/+G) apically 40µl/hour for 4 hours a day and continuously with antibiotics free advanced differentiation medium supplemented with relevant hormones basally.

This intermittent flow regime was repeated daily, the exact volume of outflows was measured, and the outflows were stored at –80°C for further analysis. The outflows collected at 72 hours were used for cytokine analysis and chip digests collected after 96 hours of co-culture were used to enumerate engrafted bacteria. To obtain chip digests, both chip channels were digested using 1mg/ml collagenase IV (gibco, 17104019) in TrypLE Express (Gibco, 12604021) for 1.5 hours or until the cells detached and the mammalian cells were counted manually using Burker Counting Chambers with Trypan Blue as described previously by our group with volume of each chip digest being monitored(*52*). Aliquots of the chip digests were stored in glycerol at – 80°C for bacterial engraftment analysis. CFUs of bacteria present on the chips at 96 hours were enumerated using spread plating the frozen chip lysates on Brucella blood agar with hemin and vitamin K1 for quantification of BVC1 (Hardy, A30) and *Lactobacilli* MRS Agar for quantification of the C6 consortium(Hardy, G117). The bacterial cultures were grown under anaerobic conditions at 37°C for 72 hours. The total volume of the chip digest for each chip was noted and the CFUs are presented as CFUs per chip.

### Mucus Imaging

To visualize mucin spatial distribution within the whole chip channel, the Emulate chips were trimmed to remove excess PDMS on both sides without interfering with the microfluidic channel and air filled channels as previously published(*19*). The chips were then perfused with the mucin-binding lectins 4μg/ml Jacalin (Fluorescein; Vector Laboratories, FL-1151-5) and 5μg/ml Wheat Germ Agglutinin (CF568 red fluor, 76221-784) in Hank’s Balanced Salt Solution, HBSS++ for 2 hours at 30μl/hour and subsequently perfused with HBSS++ for additional 2 hours prior to imaging. The chips were then removed from pods and the inlets and outlets of the chips were sealed with pipette tips. The trimmed chips were placed on glass slides sideways and imaged using an Revolve (ECHO) microscope in the inverted mode.

The side-view imaging of mucus and bacteria was performed as follows. Bacterial consortia were stained with 1x BactoView^TM^ Live Green (FITC; Biotium, 40102) and thoroughly washed prior to chip inoculation. WGA was added to the apical chip perfusion medium (HBSS++) and was perfused through the chips 4hours a day at 40μl/h. The chips were visualized using the Nikon Eclipse Ti2 microscope equipped with the Andor’s Zyla sCMOS camera. To assess the effect of inflammatory cytokines on mucus production, live chips were removed from Pods and exposed apically to 150μl of the lectin Ulex Europaeus Agglutinin I (UEA I), Rhodamine (Vector Laboratories, RL-1062-2) resuspended in HBSS++ at 10μg/ml (1:200 dilution) for 20 minutes at 37°C by gravity flow followed by two gravity washes with 150μl HBSS++. The chips were visualized in a standard top view mode using the Revolve (ECHO) Microscope.

### Effect of BVC1 on fibroblasts, PBMCs and neutrophils in conventional culture

Primary endocervical stromal cells were seeded into 96 well plates with 10,000 cells per well and allowed to adhere overnight. The next day, the adherent fibroblasts were washed 2x with PBS-/– and antibiotic free DMEM supplement with the fibroblast growth kit-low serum (ATCC, PCS-201-041) and BVC1 was added at multiplicity of infection 5 (MOI5). Finally, either PBMCS or neutrophils derived freshly from whole blood in vacutainer derived from a single donor (Research Blood Components, LLC) were added to relevant experimental groups 7 hours after the bacteria were added. The immune cells were added at 150 000 cells per well. The conditioned media were collected the next day (23 hours after the addition of immune cells) and analyzed for MMPs and cytokines.

### PBMC and Neutrophil isolation from whole blood

For immune cell isolation, 15 mL of blood (Research Blood Components, LLC) was diluted by gently adding 15 mL of PBS in a 50 mL tube and incubated with 7.5 mL of 6% dextran (Sigma Aldrich) to allow red blood cell sedimentation. The upper phase was then collected and gently overlaid onto 10 mL of Ficoll Paque Plus (Ge Healthcare), and the tube was centrifuged at room temperature at 400 x g for 30 minutes, with acceleration and brake set at the minimum allowed by the centrifuge. The PBMCs, contained in the ring between the Ficoll and the plasma, were collected, placed into a new 50 mL tube, washed by filling the tube with PBS, and centrifuged at 4°C at 300 x g for 10 minutes, with acceleration and brake set at the minimum allowed by the centrifuge.

For neutrophil isolation, the pellet sedimented after the Ficoll centrifugation was subjected to osmotic lysis with 18 mL of ddH_2_O for 10 seconds to lyse red blood cells, and then the osmolarity was restored by adding 2 mL of PBS 10X. As with the PBMCs, neutrophils were also washed by filling the tube with PBS and centrifuged at 4°C at 300 x g for 10 minutes, with acceleration and brake set at the minimum allowed by the centrifuge. The cells were then counted and resuspended in the culture medium.

### Cytokine and MMP multiplex analysis

The secreted cytokines and MMP levels were evaluated using custom R&D Luminex kits. The analyte concentrations were measured using a Bio-Plex 3D suspension array system and analyzed with Bio-Plex Manager software (Bio-Rad, v 6.0). The cytokine panel consisted of Il-1α, IL-1β, TNF-α, IL-6, IL-8 and CXCL-10, the MMP panel consisted of MMP-1, MMP-2, MMP-8, MMP-9, MMP-12, MMP-13, Cathepsin S and also included Pro-Collagen 1 alpha 1 (Human Luminex® Discovery Assay, R&D Systems). Apical chip cytokines were collected through perfusing the chips at 40μl/hour for 4 hours once every 24 hours. As the total apical outflow volume varies slightly from chip to chip, the total collected volume per chip was measured and the Luminex concentration readout was multiplied by the chip volume to obtain picogram levels of given analyte per chip per day and then divided by the number 41178 which was determined to be an average number of endocervical epithelial cells in the apical channel; and finally multiplied by 100 000 to obtain the value pg/10^5^ cells. Basal channel is perfused continuously and the basal channel cytokines levels are presented as pg/ml or as fold change compared to relevant controls.

### Microscopic Imaging

Phase contrast images of live chips were obtained using the Revolve (ECHO) microscope in an inverted mode using a custom chip holder. Bacteria on the chip were visualized using a DM IL LED inverted laboratory microscope equipped with the LEICA DFC400 camera. For immunofluorescence microscopic imaging, chips and cells were fixed using 4% paraformaldehyde (VWR, BT140770-10X10, PBS diluted), washed 3x with PBS and permeabilized with 0.1% Triton^TM^X-100 (Sigma, X100-100ML) in PBS for 15-20 minutes. The chips/cells were subsequently blocked with 1% Bovine Serum Albumin (Sigma Aldrich, A9418) in PBS for 45 minutes. Primary antibodies were diluted in blocking solution and allowed to bind to their targets overnight at 4°C. Following a wash step with PBS, the samples were incubated with secondary antibodies (1:1000) and Phalloidin to visualize the F-actin for 2 hours. Finally, the nuclei were stained with Hoechst 33342 for 10 minutes. The list of antibodies and products used in this study is listed in the **Supplementary Table 4**. Fixed chips and 2D cultured cells were visualized using an inverted microscope (Axio Observer Z1; Zeiss). Live chips stained with UEA1 and PBMCs were visualized using the Revolve (ECHO) microscope in an inverted mode using a custom chip holder.

Collagen levels in control groups and cytokine treated groups were evaluated on day 3 of treatment and day 14 of total culture using second harmonic imaging using Leica SP5 and the Leica Stellaris confocal microscopes equipped with coherent Chameleon Vision II. The chips were placed on a cover glass (VWR micro cover glass, 24 x 50mm cat. No 48382-136) and imaged using 25x water objective. The imaging plane was chosen to correspond to the fibroblast layer adjacent to the chip porous membrane. Multiphoton excitation was used to generate second harmonic images with the non-descanned detector filters with BP 430-480 and BP 500-550 laser tuned to 900nm. Three chips per group were analyzed with multiple independent spots per group.

### Gene expression analysis

Cells were harvested from apical and basal channels of the Endocervix chip using Buffer RLT plus (Qiagen, 1053393). Lysates were stored at –80°C prior to processing. Total RNA was isolated from cell lysates using RNeasy plus micro kit (Qiagen, 74034) according to manufacturer’s instructions. RNA was quantified and quality was assessed via NanoDrop One UV-Vis Spectrophotometer (Thermo Fisher). Complementary DNA (cDNA) was synthesized using the Omniscript RT (reverse transcription) kit (Qiagen, 205111) with a mix of OligodT(15) primers (Promega, C1101) and Random primers (Promega, C1181). Each cDNA synthesis reaction was prepared using 50 ng – 1 ug of RNA depending on RNA yield. Following cDNA synthesis, qPCR was performed using SsoAdvanced Universal SYBR® Green Supermix (Biorad, 1725272) on CFX96 Touch Real-Time PCR Detection System (Biorad). The thermocycling protocol was as follows, initial hold ((cDNA denaturation/polymerase activation) at 95°C for 2 minutes 30 seconds), followed by 40 cycles of denaturation (95°C for 15 seconds) and annealing/extension (60°C for 25 seconds) with plate reading, followed by melt curve. Relative RNA levels were quantified using the ΔΔCt method for (*COL1A1*, *COL3A1*, *PGR*, *PTGS2*) or Pfaffl method (*MUC5AC*, *MUC5B*) and normalized to *GAPDH*.

Primers for use in qPCR were designed for target genes using NCBI Primer BLAST. Primers were designed to span exon-exon junctions, to be 18-22 bp in length, GC content between 40-60%, primer melting temperature between 57 and 63 °C, PCR product length of 100-160. All primer sets were ordered from IDT (**Supplementary Table 5**). Melt curve analysis and PCR product gel electrophoresis confirmed successful and specific amplification of targets. Standard curves for each primer set were generated and primer efficiencies were calculated.

Total epithelial (apical channel) and stromal (basal chip channel) RNA was collected using RLT buffer (Qiagen). Total RNA or single cells in freezing media were submitted to Genewiz (Azenta Life Sciences) for next generation sequencing (Standard RNA-seq, polyA selection with ERCC spike-in, sequencing using an Illumina HiSeq for 2×150bp, ∼350M PE reads, single index). RNA-seq pipeline was conducted in Pluto where paired_end FASTQ files were processed using the nf-core-rnaseq pipeline (v3.12.0)1,2 (auto strandedness). Adapter sequences were removed withTrim Galore3. Reads were aligned with STAR4 to GRCh38 (NCBI, p.14, release 110) and quantified to gene counts using RSEM5. Analysis and figures for transcriptomic analyses were generated and analyzed using Pluto (https://pluto.bio).

For differential expression analysis, genes were filtered to include only genes with at least 3 reads counted in at least 20% of samples in any group. Differential expression analysis was then performed with the DESeq2 R package1, which tests for differential expression based on a model using the negative binomial distribution. Note: differential expression results shown here do not account for any experimental covariates that may be present (e.g. batch, paired subjects, etc).Log2 fold change was calculated for the comparison.

Gene set enrichment analysis (GSEA) was using the fgsea R package and the fgseaMultilevel() function 1. The log2 fold change from the differential expression comparisons was used to rank genes. C5: Gene Ontology gene sets – biological process gene set collection from the Molecular Signatures Database (MSigDB)2,3 was curated using the msigdbr R package4. Prior to running GSEA, the list of gene sets was filtered to include only gene sets with between 5 and 1000 genes. The false discovery rate (FDR) method was applied for multiple testing correction2. FDR-adjusted p-values are shown on the y-axis of the volcano plot. An adjusted p-value of 0.05 was used as the threshold for statistical significance.

### Proteomic Analysis

For secretome analysis, standard chip effluents were used obtained by perfusing HBSS++ for 4 hours a day at the rate 40μl/hour. As our analyses revealed mucus accumulates on the chip despite the period perfusion regime, for mucus plug analysis, the mucus was collected by incubating the apical chip content with 50 μL of 20mM N-acetyl-cysteine (NAC) for 3 hours at 37°C as previously described. The frozen chip outflows were thawed at room temperature and allowed to react individually (1:1 v:v) with 30μL of 6M guanidinium chloride solution for 1 minute under vortex. The samples were then loaded to 10KDa PALL filters and centrifuged at 4000g for 5 minutes on Eppendorf Centrifuge 5420. For reduction and alkylation, 10μL of 10mM TCEP was added to each filter for reduction at 350 rpm and 25°C for 45 minutes on Eppendorf Thermo Mixer C. All filters were centrifuged at 4000g for 5 minutes for cleanup, then 10μL of 10mM Iodoacetamide (IAA) was added to the reduced samples for alkylation at 350 rpm, 25°C for 15 minutes. Trypsin Platinum protease MS grade from Promega (100μg/mL) was used for the digestion of the samples. 100μL (0.1μg) of stock trypsin solution in 50mM Triethylammonium bicarbonate (TEAB) in water (pH 8.5) was added directly to the top of each S-Trap mini column using a 1:50 ratio (Trypsin:Protein) and samples were incubated for 2 hours. After digestion was completed 80μL of 50mM TEAB was added directly to each filter and then centrifuged at 4000g for 2 minutes. Samples, now peptides, were labeled using 10ul of TMTpro mass tag, followed by 350 rpm incubation at room temperature for 45 minutes, ensuring covalent binding between tags and peptides. Labeling reaction was quenched using 3μL of 5% hydroxylamine, then samples were pooled into one 2mL Eppendorf tube and dried using Eppendorf Vacufuge plus. Dried samples were resuspended in 120μL of 25 mM TEAB and vortexed to ensure full solubility. Samples were then fractionated using Pierce^TM^ High pH Reversed-Phase Peptide Fractionation Kit Columns (Thermo Scientific). Each fraction contained 120μL of volume for a total of 10 different fractions. Each fraction was centrifuged at 3000g for 2minutes, flowthroughs were collected and transferred to HPLC vials. All fractions were dried and resuspended in 6μL of 0.1% Formic Acid in ultrapure HPLC grade water and were then injected into LC-MS/MS.

After separation each fraction was submitted for a single LC-MS/MS experiment that was performed on a Exploris 240 Orbitrap (Thermo Scientific) equipped with NEO (Thermo Scientific) nanoHPLC pump. Peptides were separated onto a 300µmx5mm Pepmap C18 trapping column (Thermo Scientific, Lithuania) followed by DNV PepMap Neo 75umx150mm analytical column (Thermo Scientific, Lithuania). Separation was achieved through applying a gradient from 5–25% ACN in 0.1% formic acid over 120 min at 250 nL/min. Electrospray ionization was enabled through applying a voltage of 2 kV using a PepSep electrode junction at the end of the analytical column and sprayed from stainless still PepSep emitter SS 30µm LJ (Bruker, MA). The Exploris Orbitrap was operated in data-dependent mode for the mass spectrometry methods. The mass spectrometry survey scan was performed in the orbitrap in the range of 450 –900 m/z at a resolution of 1.2 × 10^5^, followed by the selection of the ten most intense ions (TOP10) ions were subjected to HCD MS2 event in the orbitrap part of the instrument. The fragment ion isolation width was set to 0.8 m/z, AGC was set to 50,000, the maximum ion time was 150 ms, normalized collision energy was set to 34V and an activation time of 1 ms for each HCD MS2 scan.

Raw MS data were submitted for analysis in Proteome Discoverer 3.1.683 (Thermo Scientific) software with Chimerys. Assignment of MS/MS spectra was performed using the Sequest HT algorithm and Chimerys (MSAID, Germany) by searching the data against a protein sequence database including all entries from the Human Uniprot database (*Homo Sapiens* Human Uniprot) and other known contaminants such as human keratins and common lab contaminants. Sequest HT searches were performed using a 20 ppm precursor ion tolerance and requiring each peptides N-/C termini to adhere with Trypsin protease specificity, while allowing up to two missed cleavages. 18-plex TMT tags on peptide N termini and lysine residues (+304.207146 Da) was set as static modifications and Carbamidomethyl on cysteine amino acids (+57.021464 Da) while methionine oxidation (+15.99492 Da) was set as variable modification. A MS2 spectra assignment false discovery rate (FDR) of 1% on protein level was achieved by applying the target-decoy database search. Filtering was performed using a Percolator (64bit version). For quantification, a 0.02 m/z window centered on the theoretical m/z value of each the 18 reporter ions and the intensity of the signal closest to the theoretical m/z value was recorded. Reporter ion intensities were exported in the result file of Proteome Discoverer 3.0 search engine as excel tables. The total signal intensity across all peptides quantified was summed for each TMT channel, and all intensity values were adjusted to account for potentially uneven TMT labeling and/or sample handling variance for each labeled channel. Upregulated and downregulated data (<30kDa) were plotted as log2FC. Targets with p value<0.05 and adjusted p value<1.6 were visualized.

Venn Diagram generation was carried out using the Ugent bioinformatics website (www: bioinformatics.psb.ugent.be/webtools/Venn/). The whole mucus content was cleaved from the apical compartment of Endocervix chips (controls, estradiol, PAH perfused) from two different donors using N-acetyl-cysteine (NAC) and the cleaved content was analyzed using mass spectrometry as described above. g:Profiler and specifically the g:Convert function was used to convert Accession Number to Name. In order to evaluate the similarity of the Endocervix-chip secretome to the native mucus plug a whole published dataset published by Vornhagen(*21*) and colleagues with values p≤0.992 was used and compared to our dataset using the Venn Diagram function available at the Ugent bioinformatics website.

### Glycomic analysis of mucus

The mucus was collected by incubating the apical chip content with 50 μL of 20 mM NAC for 3 hours at 37°C as previously described(*19*). *N-glycan release* Chip mucus effluent was processed similar to previous method(*19*), samples were filtered through a Amicon Ultra-0.5 10kDa centrifugal filter (MilliporeSigma,MA) to remove salts. Dithiothreitol (DTT) in 100mM NH_4_HCO_3_, was added to the purified samples at a final concentration of 5mM. The proteins in the samples were denatured through heating in a boiling water bath for 3 minutes, alternating between the hot bath for thirty seconds and room temperature for ten seconds. At room temperature 2 units of PNGase F (New England Biolabs, MA) were added and the samples were left in a 37°C water bath overnight to cleave all N-glycans. After incubation, nanopure water was added to increase the volume of supernatant and ultracentrifuged at 200,000 x g for 45 minutes at 4°C. This separated the released N-glycans from the pellet containing the O-glycosylated protein used for *O*-glycan release. The supernatant was desalted through porous graphitized carbon (PGC) solid phase extraction (SPE) plates (Thermo, CA). The plates were activated by 80% (v/v) acetonitrile (ACN) 0.1% (v/v) trifluoroacetic acid (TFA), equilibrated with nanopure water before samples were loaded on, washed with nanopure water, and eluted with 40% (v/v) ACN 0.05% (v/v) TFA. Eluted samples were dried in vacuo and reconstituted in nanopure water for LC-MS/MS analysis.

*O-glycan release* The O-glycosylated protein pellets were frozen at –80°C, thawed, and reconstituted in 90 μL of nanopure water. Samples were sonicated for 20 minutes to facilitate dissolution of the partially insoluble denatured protein. O-glycans were released by reductive β-elimination. Proteins were treated with 10 μL of 2 M NaOH, followed by 100 μL of 2 M NaBH_4_. The samples were incubated in a 45°C water bath overnight. The reaction was then quenched over ice with the addition of 10% acetic acid until pH of 4-6 was achieved. The resulting supernatant was desalted using PGC-SPE plates, as described for N-glycans, and dried in vacuo. O-glycans were further purified and enriched using iSPE-HILIC cartridges (Nest Group). PGC eluates were reconstituted in 90% (v/v) ACN 1% TFA, loaded onto iSPE-HILIC cartridges activated with ACN 5 volume washes in 90% (v/v) ACN 1% (v/v) TFA. The samples were eluted with nanopure water 0.1%(v/v) TFA, dried in vacuo, and reconstituted in water for LC-MS/MS analysis.

*nanoLC-MS/MS* Glycomic analysis was performed on the Agilent 1200 series HPLC-Chip system using a PCG-Chip(II) coupled to an Agilent Accurate-Mass Q-TOF MS (Agilent, CA) was previously reported(*53*). Glycan isomers were chromatographically separated and introduced to the MS through the PGC microfluidic chip consisting of an enrichment and analytical column with a nano electrospray tip. Capillary voltage was adjusted between 1850-2000 V for N-glycans but lowered to 1650-1850 V for O-glycans to maintain a stable electrospray. MS spectra were acquired in positive ionization mode, with m/z 400-2000 for O-glycans and m/z 600-2000 for N-glycans. Fragmentation for both methods used collision induced dissociation (CID) with nitrogen gas. Raw data were processed on MassHunter Qualitative Analysis B.08 software (Agilent, CA) with in-house imported libraries for N and O-glycans through the Find by Molecular Feature method with a mass tolerance of 10 ppm for O-glycans and 20ppm for N-glycans. The structures were manually confirmed through tandem MS and fragmentation patterns. The relative abundances were calculated by combining all isomers peaks for each structure and normalizing it to the total peak area.

## SUPPLEMENTARY FIGURES

**fig S1.**
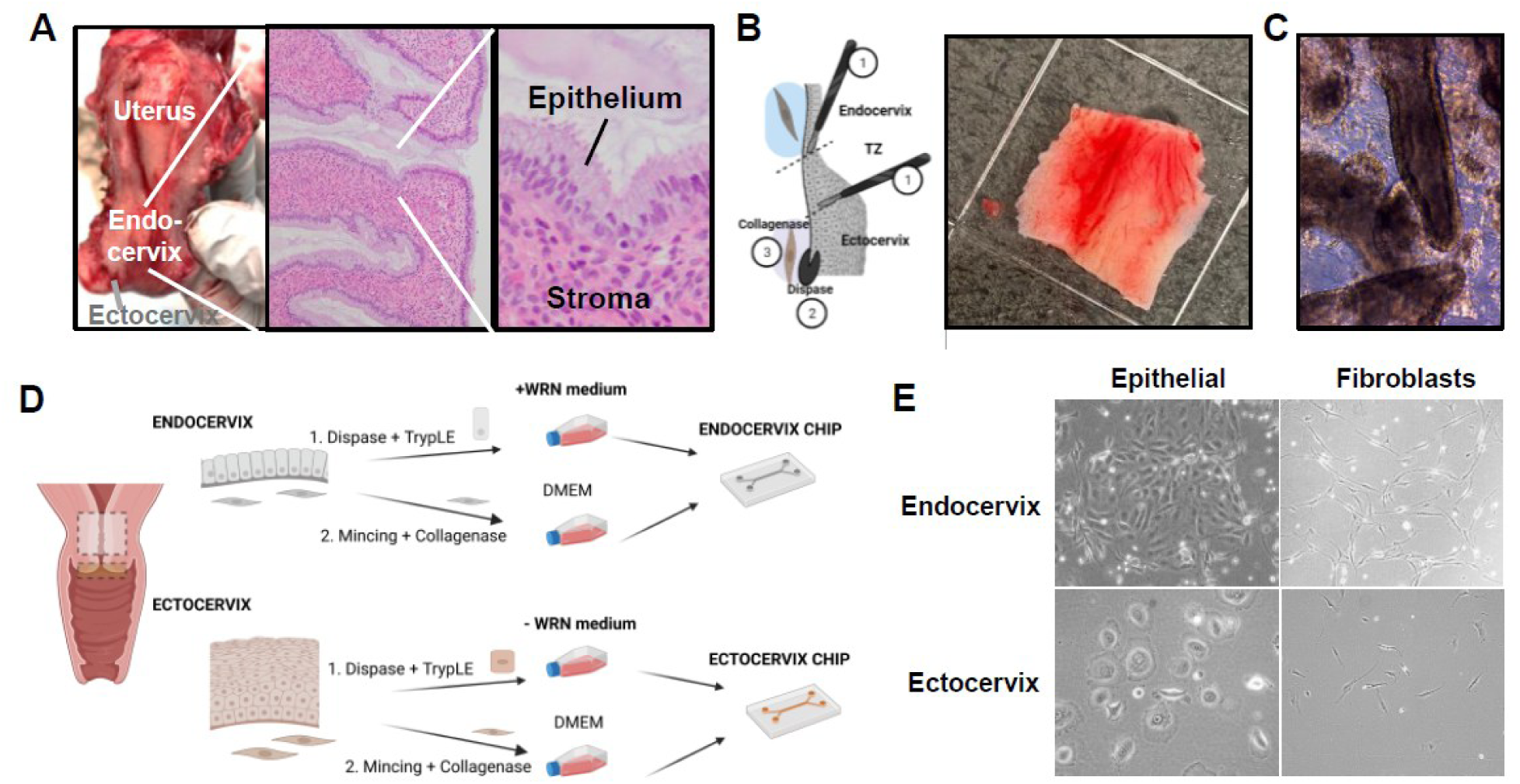
Endocervix and Ectocervix Epithelia and Stroma Isolation. **A)** Photograph and histological sections of the uterus and uterine endocervix glands. **B)** Diagram (left) illustrating the workflow for isolating the endocervix from the ectocervix whereby (1) the ectocervix and endocervix tissues are isolated by experienced pathologists at Mass General Hospital and the transition zone (TZ) is discarded. (2) The basement of both the endocervix and ectocervix is enzymatically digested using dispase and (3) collagenous stroma is dissolved using collagenase I & II. The photograph (right) shows an excised endocervix. **C)** Brightfield image of freshly isolated endocervical glands. **D)** Pictogram (created in: BioRender) outlining the isolation of endocervical and ectocervical epithelial cells (grey columnar top, brown stratified bottom) and their expansion in (+) or without (-) WRN medium (WRN=Wnt3a, R-spondin3 and Noggin). **E)** Representative images of passage 0 (P0) endocervix and ectocervix epithelia and fibroblasts in T75 flasks.

**fig S2.**
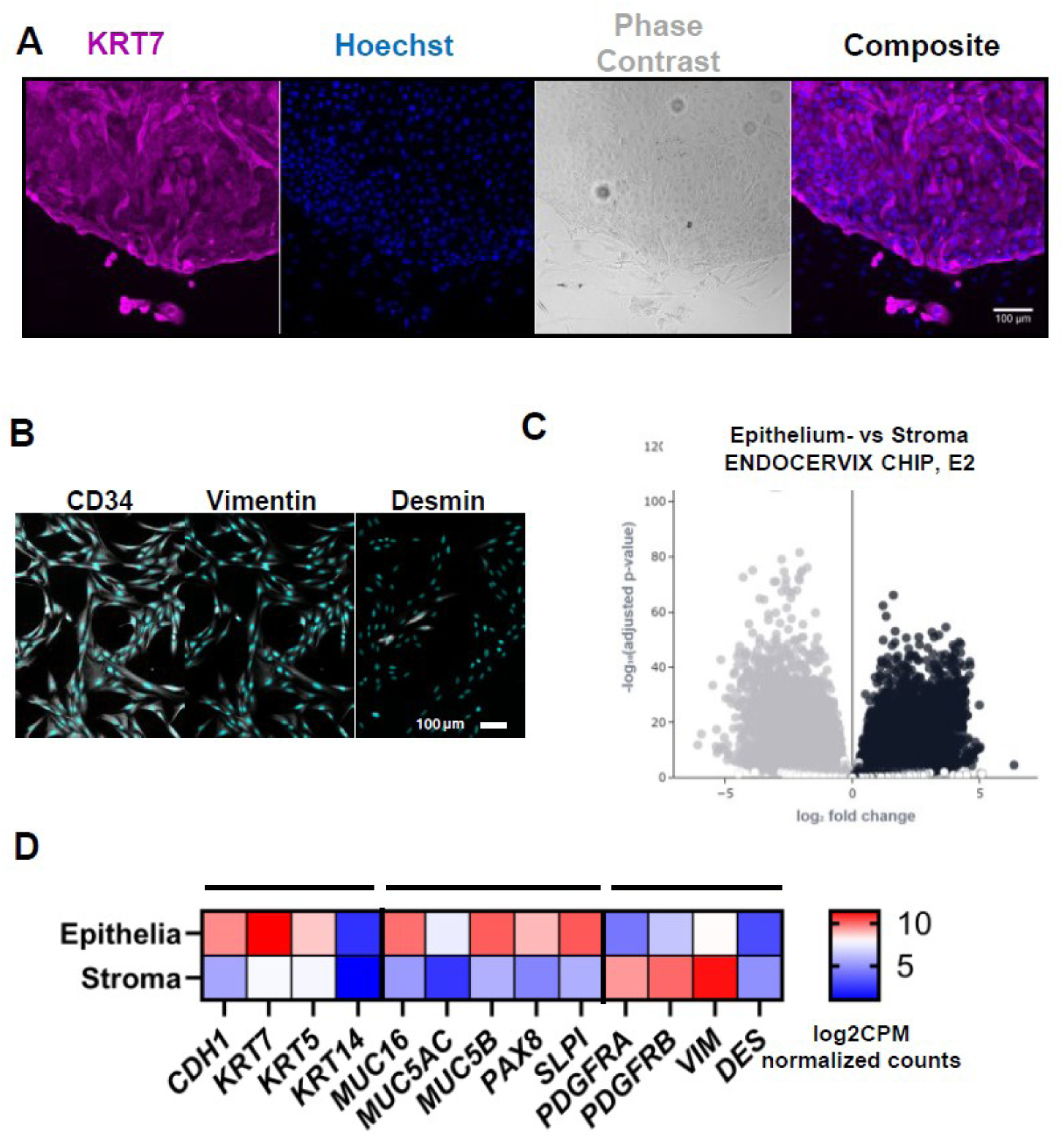
Evaluation of Endocervical Cell Identity. **A)** Immunohistochemistry image showing isolated endocervical cells are KRT7 (purple) positive. Images also show epithelial nuclei (blue), (bar, 100 µm). **B)** Immunocytochemistry images of primary endocervical stromal cells stained for CD34 (white, left), vimentin (white, middle) and desmin (white, right); and nuclei (cyan), (bar,100 µm). **C)** Volcano plot displaying log2 fold change (x-axis) vs. –log10(adjusted p-value) (y-axis) for gene expression in endocervix chip epithelia (black dots) vs. stroma (grey dots) under estradiol perfusion. (N=10-11 chips per group). **D)** Heatmap of log2CPM-normalized counts for endocervical epithelial (*CDH1*, *KRT7*, *MUC16*, *MUC5B*, *MUC5AC*, PAX8, *SLPI*), ectocervical epithelial (*KRT14*), endocervical fibroblast (*PDGRA*, *PDGFRB*, *VIM*), and smooth muscle (*DES*) genes in differentiated primary endocervix chips (N=2 donors, N=3 chips per donor).

**fig S3.**
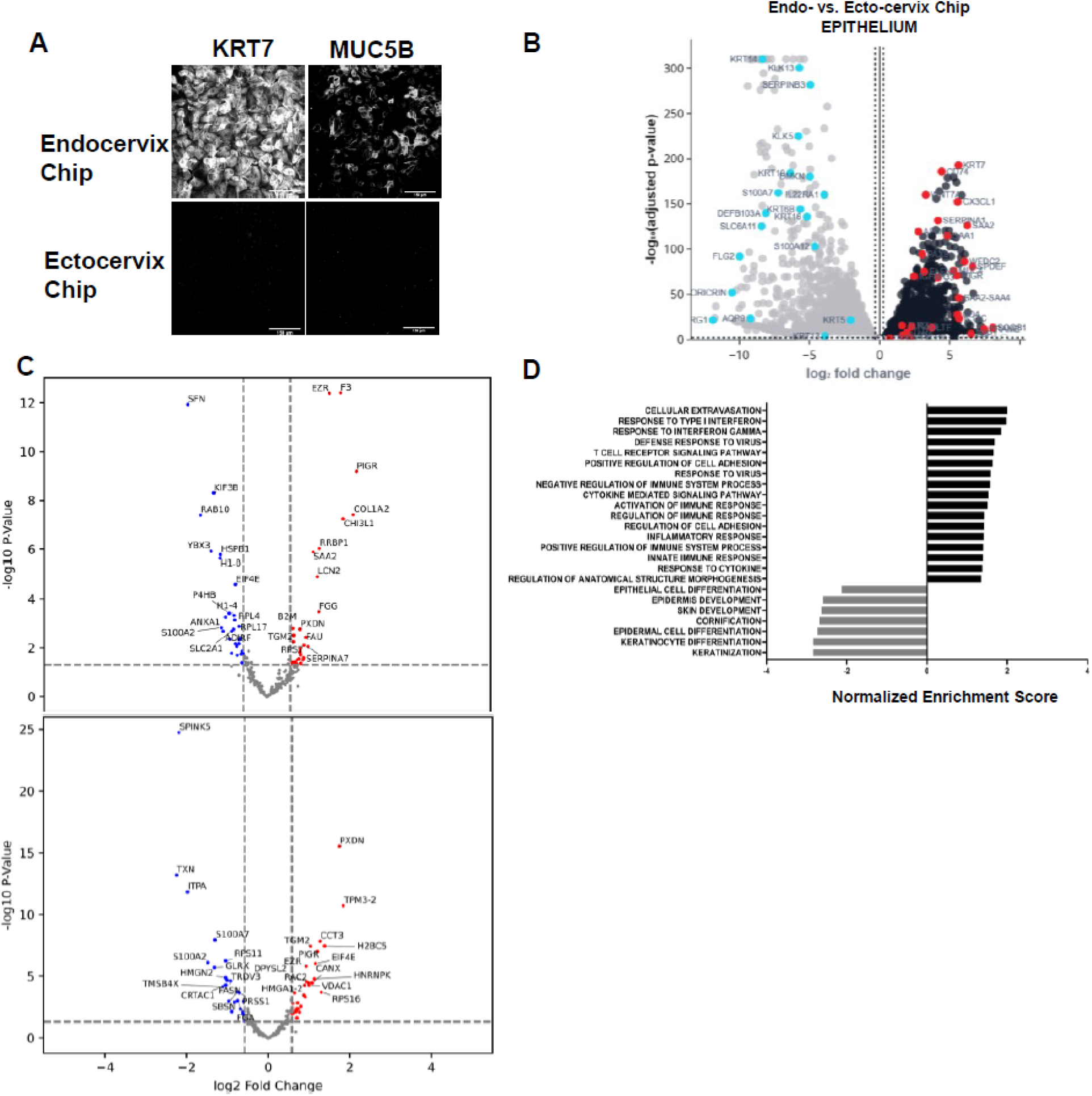
Distinct Mucus and Collagen Expression in Mature Endocervix and Ectocervix Chips. **A)** Immunofluorescence images showing differentiated Endocervix and Ectocervix Chip epithelia stained for KRT7 (white, left) and MUC5B (white, right), (bar, 150 µm). **B)** Volcano plot comparing the epithelia of donor matched Endocervix (black) and Ectocervix (grey) Chips showing the log2 fold change of each gene on the x-axis and the –log10(adjusted p-value) on the y-axis. Points represent gene with adjusted p-value ≤ 0.01 and a fold change that is either < 0.8 or >1.2. (N=4 chips per group).**C)** Volcano plots of differentially expressed proteins in the secretomes of Endocervix and Ectocervix Chips perfused with estradiol (E2). Data shown separately for two distinct donors (N=4 chips per donor) **D)** Bar plot showing the GSEA results for Ectocervix and Endocervix Chip epithelia shown in **B** for each tested gene set with a p-value ≤ 0.01. The x-axis shows the normalized enrichment score (NES), which represents the magnitude of enrichment as well as the direction (gene sets are ordered along the y-axis by NES value). (N=4 chips).

**fig S4.**
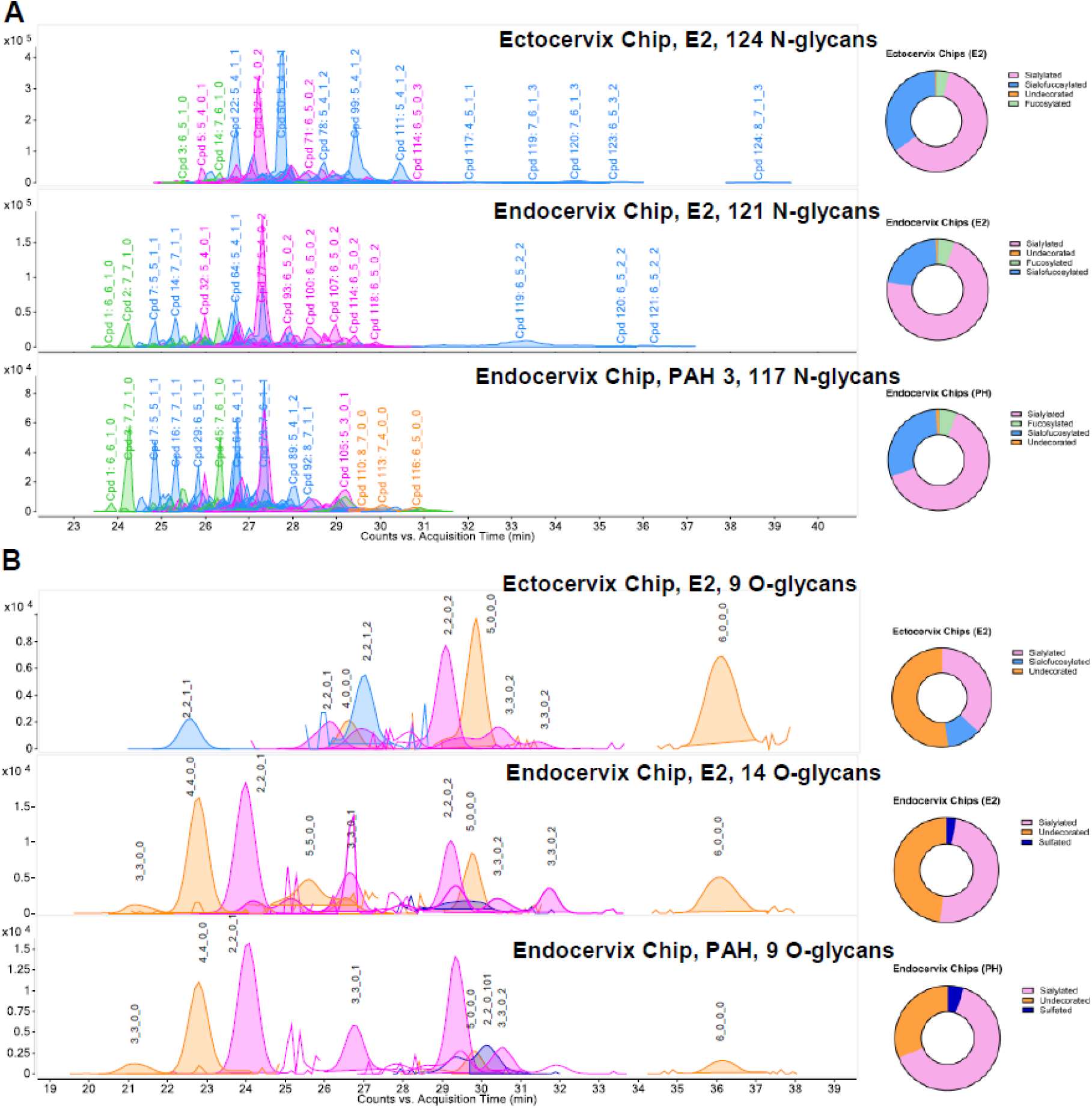
Glycomic Analysis of Endocervix and Ectocervix Chips. **A**) N-glycan profiles of NAC-cleaved apical mucus from (N=3-4 Ectocervix and Endocervix Chips). **B**) O-glycan profiles of NAC-cleaved apical mucus from (N=3-4 Ectocervix and Endocervix Chips, E2=estradiol, PAH=pregnancy associated hormones; peaks are color coded as follows: sialylated = pink, sialofucosylated = blue, undecorated = orange, fucosylated = green, sulfated = dark blue). Y-axis represents counts, X-axis represents Acquisition time (minutes).

**fig S5.**
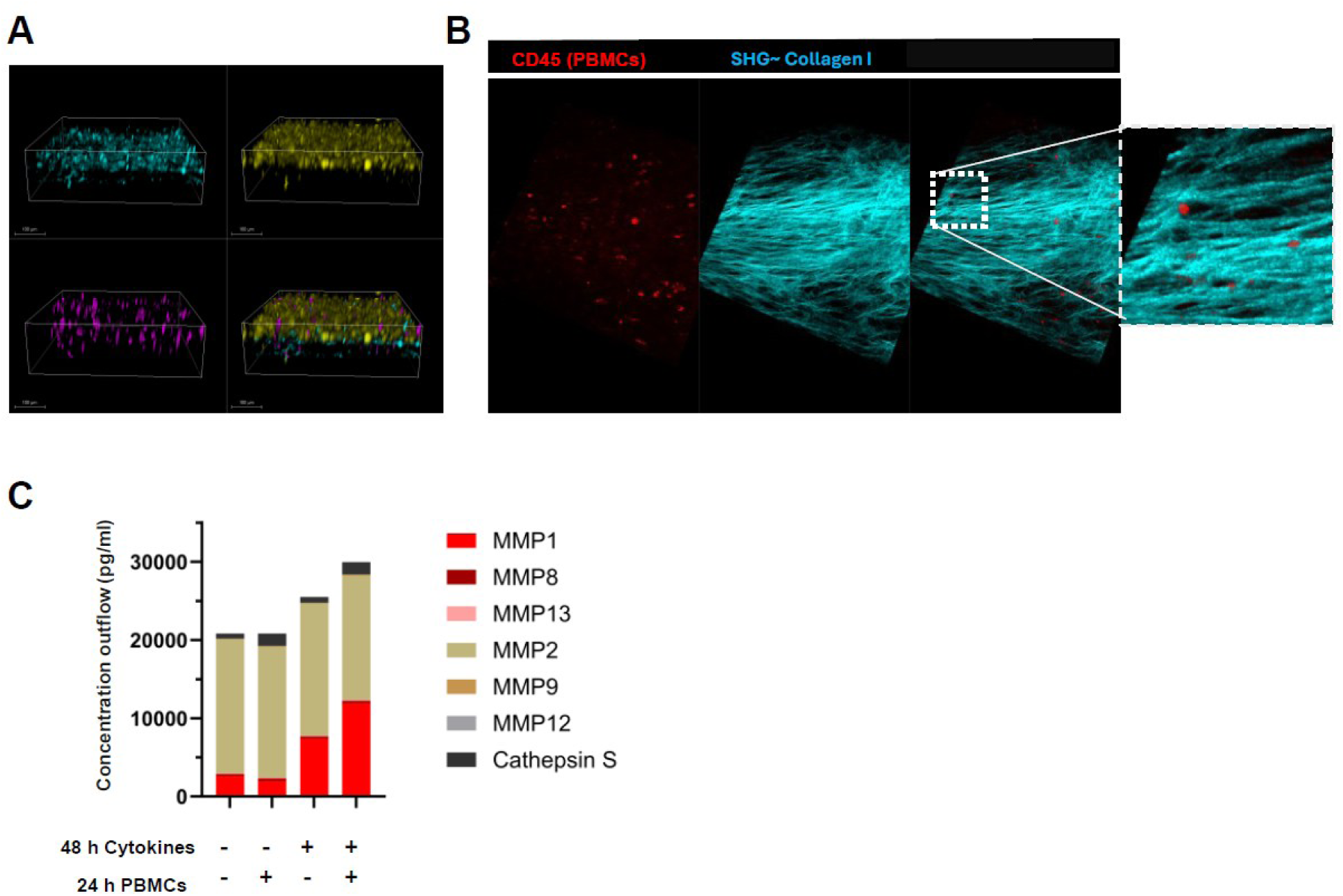
PBMC Migration in Endocervix Chips. **A**) A 3D confocal reconstruction of an Endocervix Chip showing PBMCs (magenta), collagen (cyan, SHG) and endocervical epithelial and stromal cells (yellow, autofluorescence) (bar, 100 µm). **B**) Visualization of CD45+ PBMCs (red) in between collagen I fibers(cyan, SHG) in the basal channel of the Endocervix Chip. **C**) A bar graph showing the concentration of MMPs (MMP-1, MMP-8, MMP-13, MMP-2, MMP-9, MMP-12) and Cathepsin S in basal outflows of PAH-perfused endocervix chips with/without inflammatory cytokines (IL-1β, TNF-α, IL-6, IL-8) and PBMCs (N=4 chips).

**fig S6.**
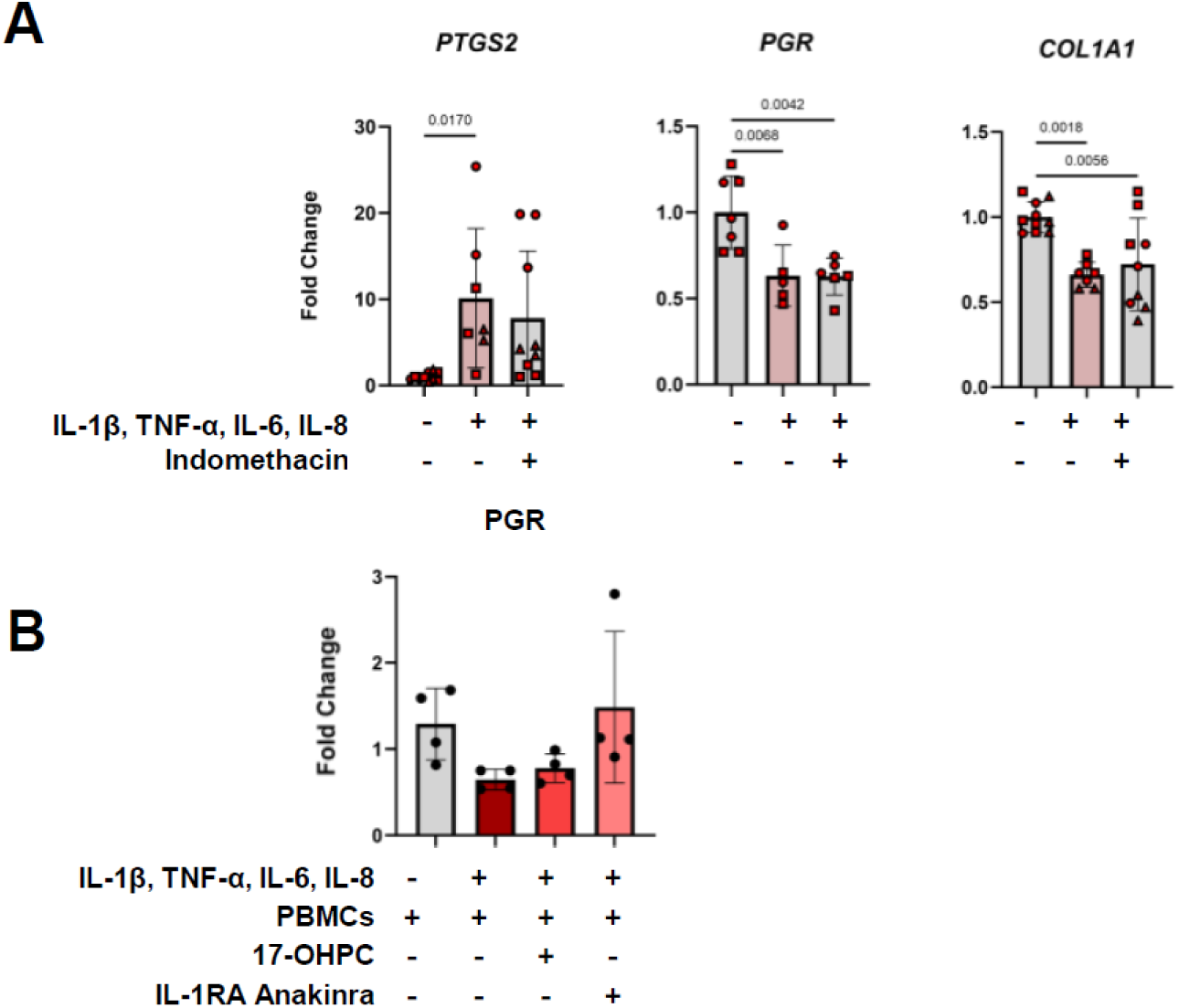
Effect of Indomethacin and IL-1RA on Endocervix Chip Stroma Under Inflammatory Conditions. **A)** Graphs showing stromal expression levels of preterm labor-associated genes (*PTGS2*, *PGR*, *COL1A1*) in Endocervix Chips pre-treated and basally perfused with Indomethacin and apically perfused with inflammatory cytokines (IL-1β, TNF-α, IL-6, IL-8). (N=7-10 chips. Mean ± SD, One-way ANOVA, expression relative to housekeeping genes and control chips (ΔΔCt)). **B)** Graph showing *PGR* expression in PAH perfused Endocervix Chips treated either with 17-OHPC or Anakinra (N=4 chips, Mean ± SD, one-way ANOVA)

**Table S1:** Hormones.

| **Estradiol (E2)** | **Company** | **Product number** | **Concentration on chips non-pregnant** |
| --- | --- | --- | --- |
| β -Estradiol (E2) | Sigma | E4389 | 5 nM |

| **Pregnancy Associated Hormones (PAH)** | **Company** | **Product number** | **Concentration on chips ~34 weeks** |
| --- | --- | --- | --- |
| β -Estradiol (E2) | Sigma | E4389 | 45.884 nM |
| Medroxyprogesterone 17-acetate (MPA) | Biovision, Sigma | B2192-100, 46412-250mg | 1 μM |
| Prolactin (PRL) hormone | R&D, PeproTech® | 682-PL, 100-07-100ug | 140 ng/ml |
| Human Chorionic Gonadotrophin (HCG) | Abcam | 51782 | 200 ng/ml |
| Human Placental Lactogen (CSH1) | Sino biologicals, Abcam | 11596-H08H, LC6FE2204, 103681-334 (EA) | 2.5 μg/ml |

**Table S2.** Endocervical Expansion WRN-Medium Composition.

| **Component** | **Final conc** | **Catalog #** |
| --- | --- | --- |
| **L-WRN Conditioned Medium** | 50 %vol/vol | Cell line: L-WRN, ATCC, CRL-3276 |
| **Advanced DMEM/F12** | - | Thermo Fisher Scientific, 12634-010 |
| **GlutaMAX** | 1x | Thermo Fisher Scientific, 35050-061 |
| **HEPES** | 12 mM | Thermo Fisher Scientific, 15630-106 |
| **B27** | 1x | Thermo Fisher Scientific, 17504044 |
| **N2** | 1x | Thermo Fisher Scientific, 17502001 |
| **Nicotinamide** | 10 mM (1.2mg/ml) | Sigma-Aldrich, N0636 |
| **NAC (N-acetyl-L-cystein)** | 1.25 mM (203.75ug/ml) | Sigma-Aldrich, A5099 |
| **Primocin®** | 100 ug/ml | InvivoGen, ant-pm-1 |
| **EGF** | 8.33 nM (50 ng/ml) | Genscript, Z00333 |
| **A8301** | 500 nM (210.79 ng/ml) | Stemcell Technologies, 72022 |
| **SB202190** | 10 uM | Stemcell Technologies, 72632 |
| **Recombinant Human FGF-10** | 100 ng/ml | Peprotech, AF-100-26-100UG |
| **Recombinant Human HGF** | 50 ng/ml | Peprotech, 100-39H-100UG |
| **Y-27632 (Dihydrochloride)** | 3.5 uM | Stem Cell Technologies, 72304 |

**Table S3:** Endocervix chip medium Expansion Stage.

| Apical Medium | Component | Content |
| --- | --- | --- |
|  | Endocervical Expansion WRN-Medium (Table S2) | 25% |
|  | Dermacult™ (StemCell Technologies) | 75% |
| Basal Medium | Component | Volume/Concentration |
|  | Basal Medium DMEM high glucose + 10% FBS | 500 ml |
|  | L-glutamine, (ATCC, PCS-201-041 kit component) | 7.5 mM |
|  | rh FGF basic, (ATCC, PCS-201-041 kit component) | 5 ng/mL |
|  | rh Insulin, (ATCC, PCS-201-041 kit component) | 5 µg/mL |
|  | Hydrocortisone, (ATCC, PCS-201-041 kit component) | 1 µg/mL |
|  | Ascorbic acid, (ATCC, PCS-201-041 kit component) | 50 µg/mL |
|  | Fetal bovine serum, (ATCC, PCS-201-041 kit component) | 2% |

**Differentiation Stage**
| Apical Medium | Component | Content |
| --- | --- | --- |
|  | HBSS++ | 100% |
| Basal Medium | Component | Volume/Concentration |
|  | Advanced DMEM/F12 (12634010) | 500 ml |
|  | L-Glutamine LifeFactor, (LifeLine, LS-1097 kit component) | 15 ml |
|  | Epinephrine LifeFactor, (LifeLine, LS-1097 kit component) | 0.5 ml |
|  | Hydrocortisone Hemisuccinate (LifeLine, LS-1097 kit component) | 0.5 ml |
|  | rh EGF LifeFactor, (LifeLine, LS-1097 kit component) | 1 ml |
|  | Extract P™ LifeFactor, (LifeLine, LS-1097 kit component) | 4ml |
|  | Apo-Transferrin Life-Factor, (LifeLine, LS-1097 kit component) | 1 ml |
|  | Rh Insulin Life Factor, (LifeLine, LS-1097 kit component) | 1 ml |
|  | HLA supplement (ATCC, PCS-201-040™ kit component) | 1.25 mL |
|  | Ascorbic acid (ATCC, PCS-201-040™ kit component) | 0.5 mL |
|  | Hormones (Supplementary Table S1) |  |

**Table S4.** List of Antibodies.

| Target | Company | Type | Product | Dilution |
| --- | --- | --- | --- | --- |
| **MUC5B** | abcam | Mouse monoclonal | ab77995 | 4µg/ml |
| **KRT7** | abcam | 555 conjugated | ab209601 | 1/250 dilution |
| **F-actin** | invitrogen | Alexa Fluor™ Plus 647 Phalloidin | A30107 | 1/400 dilution |
| **Nucleus** | vwr | Hoechst 33342 | PI62249 | 1/100 dilution |
| **CD34** | abcam | mouse monoclonal | ab8536 | 1/100 dilution |
| **Desmin** | abcam | mouse monoclonal | ab8470 | 1/100 dilution |
| **Vimentin** | abcam | 647 conjugated | ab195878 | 1/200 dilution |
| **CD45** | invitrogen | Mouse monoclonal | 14-0459-82 | 1/100 dilution |

**Table S5:**
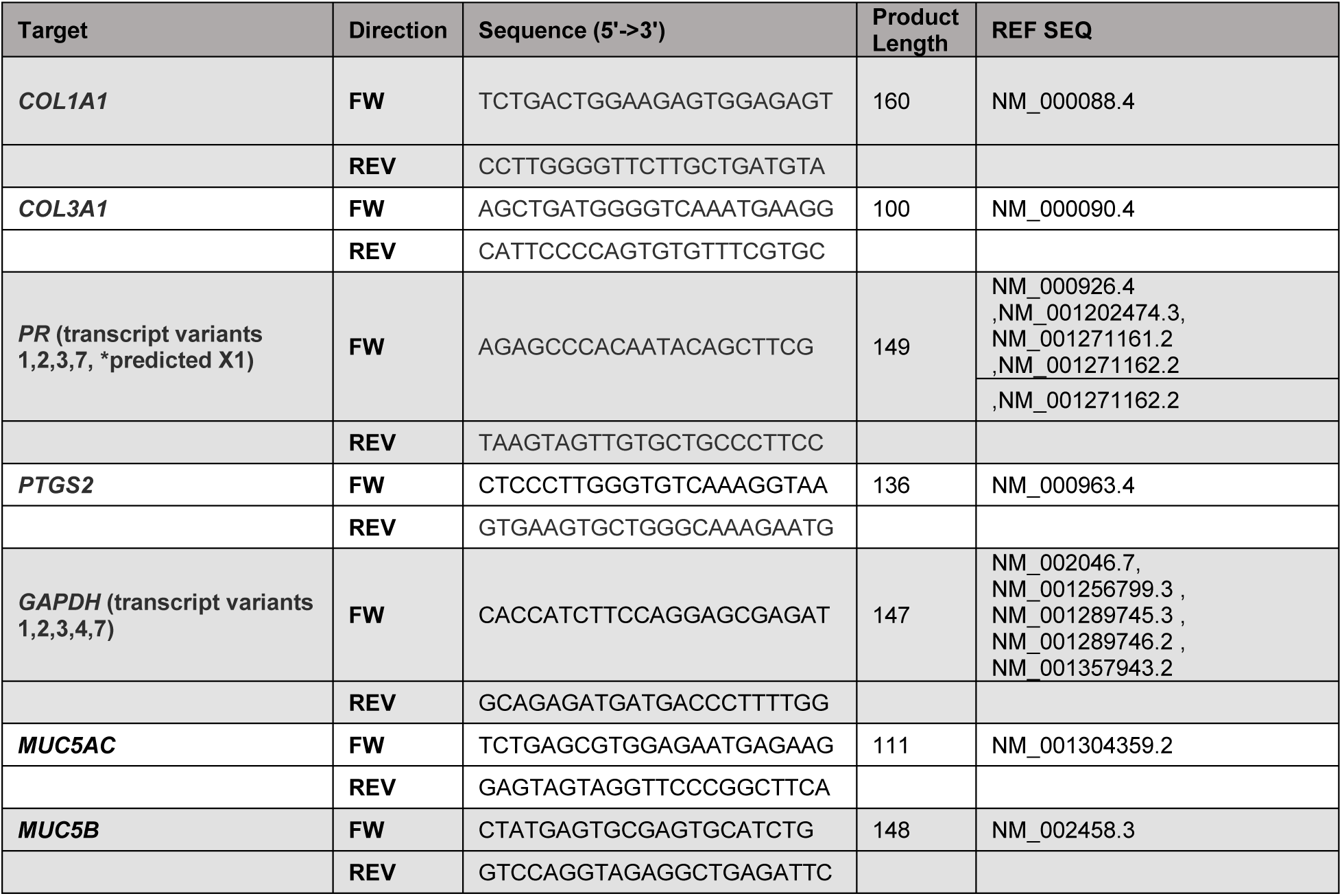
List of Primers.

